# Metapopulation model of phage therapy of an acute *Pseudomonas aeruginosa* lung infection

**DOI:** 10.1101/2024.01.31.578251

**Authors:** Rogelio A. Rodriguez-Gonzalez, Quentin Balacheff, Laurent Debarbieux, Jacopo Marchi, Joshua S. Weitz

## Abstract

Infections caused by multi-drug resistant (MDR) pathogenic bacteria are a global health threat. Phage therapy, which uses phage to kill bacterial pathogens, is increasingly used to treat patients infected by MDR bacteria. However, the therapeutic outcome of phage therapy may be limited by the emergence of phage resistance during treatment and/or by physical constraints that impede phage-bacteria interactions *in vivo*. In this work, we evaluate the role of lung spatial structure on the efficacy of phage therapy for *Pseudomonas aeruginosa* infection. To do so, we developed a spatially structured metapopulation network model based on the geometry of the bronchial tree, and included the emergence of phage-resistant bacterial mutants and host innate immune responses. We model the ecological interactions between bacteria, phage, and the host innate immune system at the airway (node) level. The model predicts the synergistic elimination of a *P. aeruginosa* infection due to the combined effects of phage and neutrophils given sufficiently active immune states and suitable phage life history traits. Moreover, the metapopulation model simulations predict that local MDR pathogens are cleared faster at distal nodes of the bronchial tree. Notably, image analysis of lung tissue time series from wild-type and lymphocyte-depleted mice (n=13) revealed a concordant, statistically significant pattern: infection intensity cleared in the bottom before the top of the lungs. Overall, the combined use of simulations and image analysis of *in vivo* experiments further supports the use of phage therapy for treating acute lung infections caused by *P. aeruginosa* while highlighting potential limits to therapy given a spatially structured environment, such as impaired innate immune responses and low phage efficacy.

## I. INTRODUCTION

The antimicrobial resistance crisis poses a major burden to global health, with 1.27 million deaths attributed to drug-resistant bacteria in 2019 alone^1^. The discovery of new antibiotics has slowed^2^, with only two new chemical classes of antibiotics approved since 2017. The antimicrobial resistance crisis has also catalyzed the search for alternative therapies, including phage therapy. Phage therapy conventionally uses bacteriophage (phage) to target and kill specific bacterial pathogens. However, the emergence of phage resistance and/or physical constraints hindering phage propagation *in situ* can limit phage treatment efficacy.

The ability of phage to kill bacteria is typically assessed through *in vitro* liquid medium assays. In these well-mixed systems, phage demonstrate efficient killing of both gram-positive^3^ and gram-negative^3,4^ bacterial pathogens. In contrast, phage are less efficient at clearing biofilms^5–8^, especially well-established and matured biofilms^5,7,8^. The spatial heterogeneity of biofilms limits phage propagation and biofilm clearance^9,10^. For instance, slow-growing bacteria living at the center of a biofilm restrict phage propagation and impede biofilm elimination^9^. Furthermore, clusters of phage-resistant bacteria can protect susceptible bacteria by blocking phage infection in mixed-strain biofilms^10^.

Applications of phage therapy *in vivo* and in the clinic require consideration of context, including spatial structure and host immune responses. For example, a three-phage cocktail targeting *E. coli* in the murine gut resulted in the coexistence of phage and bacteria^11^. This study also found a significantly higher relative abundance of phage in the luminal part of the ileum than in the mucosal part^11^, suggesting intestinal mucosa serves as a spatial refuge for bacteria. In a similar study, high T7 phage and *E. coli* titers coexist in the mice colon for up to three weeks^12^.

With 80% of isolated *E. coli* colonies susceptible to parental phage, the study proposes that susceptible bacteria in the colon may be physically protected from phage infection^12^. The host immune responses may play a significant role in shaping phage therapy outcomes *in vivo*. For instance, the synergistic interactions between phage and neutrophils were responsible for the clearance of *P. aeruginosa* lung infection in WT mice^13^. In contrast, phage therapy failed to eradicate *P. aeruginosa* infection in neutropenic mice^13,14^. Moreover, phage immunogenicity assays indicate that phage do not significantly increase cytokine production and are well-tolerated by mammalian hosts^13,15^. However, prolonged treatment (∼6 months) with intravenously administered phage might induce neutralizing antibody responses against phage^16^.

Computational models and theory provide a route to evaluate the impacts of immune responses on the potential efficacy of phage therapy. For example, pioneering modeling work integrating immune mechanisms into phage therapy^17,18^ allowed the assessment of the combined effects of phage life history traits and immune response intensity on controlling a bacterial population. However, those models allowed the immune response to grow unconstrained and assumed bacteria cannot evade the immune responses, leading to scenarios where the host immune system alone could control bacterial infection^17,18^. Recent work incorporating more nuanced models reveals the necessity of both phage and a sufficiently competent immune system to clear the infection^13,19^. Such synergy may also apply to phage-antibiotic combination therapy^20^. Additionally, models integrating adaptive and innate immune responses against phage predict that, if active, immune effects could contribute to phage therapy failure in controlling bacterial infection^21^. It is important to note that the discussed models lack an explicit representation of space, a critical component when modeling phage therapy *in vivo*.

Spatially explicit models are suitable tools for assessing the impact of the spatial organization of phage and bacteria on infection dynamics and therapeutic outcomes. For instance, individual-based models (IBMs) of phage therapy identify that spatial structure limits phage dispersion and enables spatial refuges for bacteria to survive phage infection, facilitating the coexistence of phage and bacteria^22,23^. In the context of phage-antibiotic combination therapy, an IBM found that double-resistant mutants emerged when phage dispersion was restricted and antibiotics were heterogeneously distributed. This led to the creation of drug-free spatial refuges where bacteria replicate and acquire phage and antibiotic-resistant mutations^22^. Moreover, an IBM of phage treatment of a mixed-strain biofilm identified that fitness costs associated with phage resistance, coupled with spatial protection of susceptible cells vs. phage infection by clusters of resistant bacteria, facilitated the long-term coexistence of susceptible and resistant strains^10^.

Spatial organization into subpopulations has been proposed as a key factor in fostering the persistence of predatorprey interactions^24^. Metapopulation models are spatially explicit models, representing environments as interconnected patches where local populations reside and interact via individuals moving among the patches. A metapopulation model of the host-parasitoid pair, *Callosobruchus chinensis* and *Anisopteromalus clalandrae*, found that the persistence of the host-parasitoid interaction increased with spatial subdivision and with the number of patches^24^. In a host plant-pathogen metapopulation model, limited host dispersion led to increased pathogen diversity while increased host dispersion homogenized the dynamics of local populations and decreased pathogen diversity^25^. Additionally, a metapopulation model exploring pathogen circulation in healthcare settings found that pathogen control via host sanitation, rather than environment sanitation, led to more rapid elimination of the circulating pathogen^26^. It is worth noting that a metapopulation modeling framework and its ecological principles can be adapted to describe microbial ecological dynamics within the lung environment, providing insights into the influence of spatial structure on pathogen persistence.

In this study, we introduce a metapopulation model that integrates lung spatial structure and host immune responses to evaluate their combined impact on phage therapy outcomes for acute pneumonia caused by *P. aeruginosa*. The model simulates ecological interactions between two bacterial strains (phage-susceptible and resistant), phage, and the innate immune response within a network structure resembling the branching pattern of the lungs. Our findings reveal that the clearance of a *P. aeruginosa* infection is contingent upon sufficiently active innate immune states and suitable phage life history traits. Moreover, we note the spatial model requires higher innate immune response levels to increase the chances of therapeutic success, in contrast to a well-mixed model, which predicts infection resolution with lower innate immune levels, highlighting the spatial structure’s role in shaping phage therapy outcomes. Throughout our simulations, we observe a spatial pattern wherein infection clears faster at the network’s bottom compared to the top, demonstrating robustness across varied distributions of bacterial inoculum and phage dose. Lastly, our analysis of *in vivo* mice lung infection data^13^ identifies a concordant and statistically significant pattern: a bottom-to-top spatial transition in pathogen clearance, aligning with the predictions of the metapopulation model.

## II. MODEL

The metapopulation network structure is based on the geometry of a symmetrical bronchial tree with a dichotomous branching pattern (Fig. 1a). The nodes of the network represent the airways, and the network links represent the branching points on the bronchial tree. We assume bacteria colonize the airways (nodes) and spread across the network through the network links. We consider the structure of the bronchial tree to be perfectly symmetrical, such that the left and right sides of the tree are identical. Hence, in the absence of stochastic effects, the dynamics at every node of the same tree generation, *g*, will be identical. Therefore, we can consider only one node in each generation without losing any information on the whole network. The final network topology consists of a chain of connected nodes (Fig. 1a-right), where each node represents one airway per generation (*g*) of the tree.

**FIG. 1:**
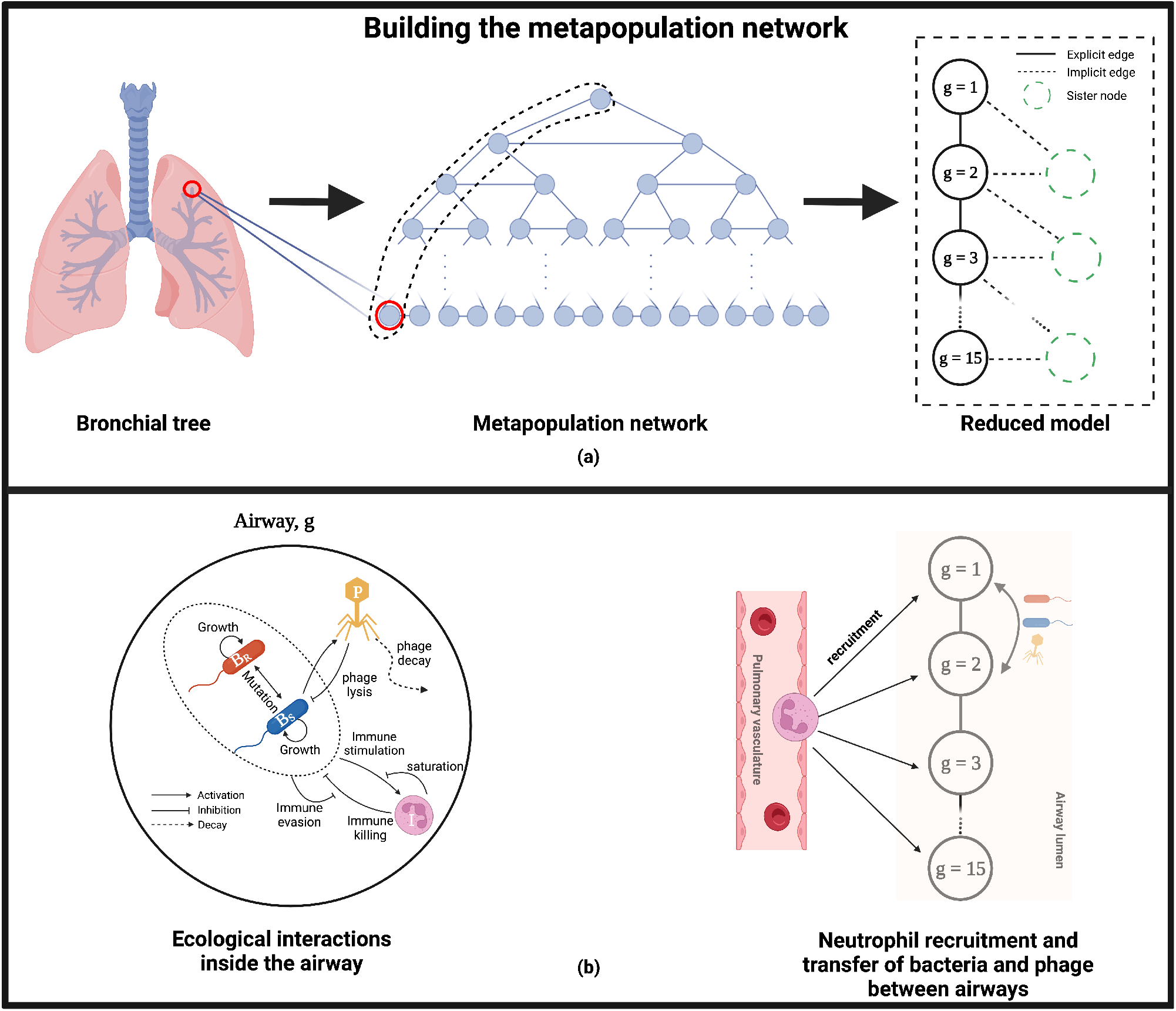
Schematic of the metapopulation network model of phage therapy of a *P. aeruginosa* infection. The structure of the metapopulation network is based on the geometry of a symmetrical bronchial tree with a dichotomous branching pattern (a). The airways of the bronchial tree (a-left) are represented by the network nodes (a-middle), while the network links represent the branching points of the bronchial tree. For example, the section that connects the trachea to the left and right main bronchi is considered a branching point. We assume that the bronchial tree is symmetrical such that the left and right parts of the tree are identical, and so are their dynamics in a deterministic system. Therefore, we only focus on one side of the tree (the dashed line on the middle network) and reduce the number of nodes in our original network to one node per generation. The final network topology of this reduced model consists of 15 connected nodes forming a chain (a-right). The degree of the network nodes is 4, except for the first and last nodes, which have a degree of 2. In panel (b), we show the ecological interactions between phage-susceptible bacteria (*B*_*S*_), phage-resistant bacteria (*B*_*R*_), phage (*P*), and the host innate immune response (*I*) at the node level (b-left). Phage infect the susceptible strain, while the resistant strain is targeted only by the innate immune response. The immune response grows in the presence of bacteria and targets both bacterial strains. Neutrophils are recruited to the site of the infection from the pulmonary vasculature, while phage and bacteria transfer between connected nodes to spread across the network (a-right).

To determine the number of generations of the metapopulation network, we estimate the volume of the network in terms of the ∼1 ml total lung capacity (TLC) of mice^27^. To do so, we calculate the volume of individual airways of different generations using the anatomical airway information^28^. Then, we consider the number of airways per generation (i.e., 2^*g*−1^ airways per generation, *g*) to estimate the network volume. We estimated that 15 generations yield a network volume of 0.9 ml that does not exceed the mouse TLC.

Nonlinear interactions between bacteria, phage, and neutrophils govern the population dynamics at the node level, Fig. 1b-left. We represent the ecological interactions between bacteria, phage, and neutrophils using a system of nonlinear, ordinary differential equations (Eqns 1-4) for each generation (*g*). We propose a model with two *P. aeruginosa* strains, one of which is phage-susceptible (*B*_*S*_) and a second which is phage-resistant (*B*_*R*_). The growth of bacteria is density-dependent and is limited by the carrying capacity, *K*_*C*_. *B*_*S*_ can be infected and lysed by phage (*P*) while *B*_*R*_ resists phage infection. Phage, *P*, replicate inside bacteria and decay in the environment at a rate *ω*. The host innate immune response, *I*, is activated by bacteria and recruited at a maximum rate *α*. Neutrophils target and kill both *B*_*S*_ and *B*_*R*_ bacterial strains.

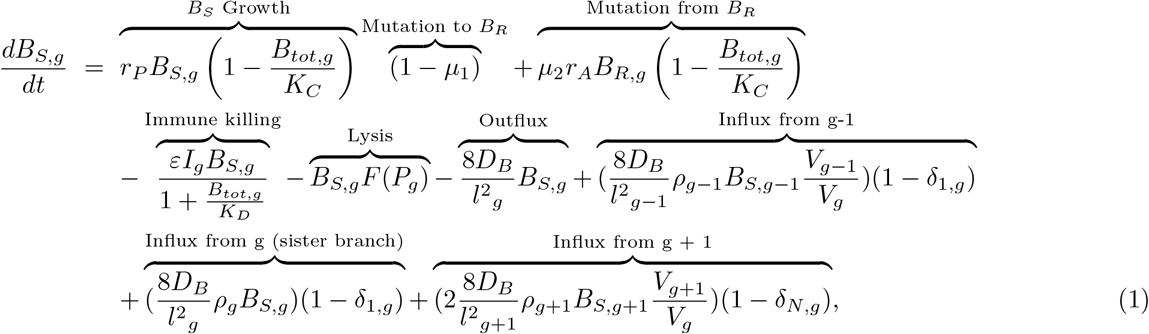

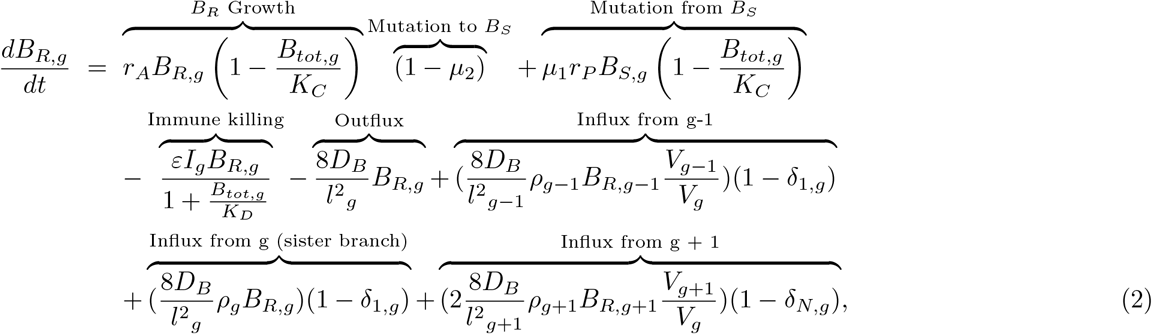

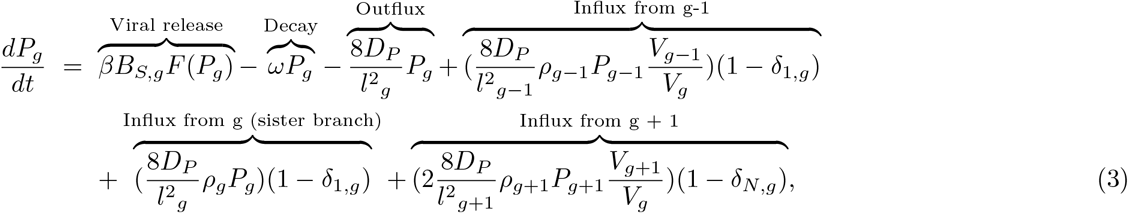

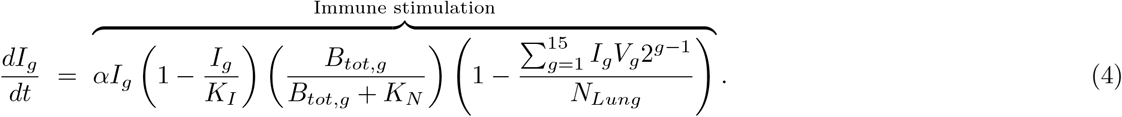

In our model, *B*_*S*_ grows at a maximum rate *r*_*P*_, while *B*_*R*_ grows at a maximum rate *r*_*A*_, and total bacterial density is represented by *B*_*tot*_ = *B*_*S*_ + *B*_*R*_. Phage infect and lyse the phage-susceptible bacteria (*B*_*S*_) at a rate *F* (*P*) with a burst size of *β*. Locally, the innate immune response (*I*) saturates at a maximum density of *K*_*I*_ . The immune response is also globally constrained by the total number of neutrophils available in the lungs, *N*_*Lung*_. *K*_*N*_ is a half-saturation constant, i.e., the bacterial density at which the growth rate of the immune response is half its maximum. Both susceptible and resistant bacteria are killed by the innate immune response with a maximum killing rate *ε*. However, at high densities, bacteria can activate mechanisms such as biofilm formation and production of virulence factors to evade the host immune response and reduce the immune killing efficiency^19^. *K*_*D*_ is the bacterial density at which the immune killing rate is half its maximum.

Bacteria and phage can hop between connected nodes (Fig. 1b-right). The rate at which they hop is determined by the time, *τ*_*g*_, required for each species to cross half of the airway length (*l*_*g*_) via diffusion *D*. The hopping rate is calculated as the reciprocal of these times,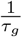. In the model, influx and outflux terms govern the transfer of bacteria and phage between neighboring nodes. We assume a homogeneous spreading between airways such that the links between nodes are non-weighted and the flux from node *j* to node *g* is inversely proportional to the degree of node *j* (i.e., *ρ*_*j*_ = 1*/d*_*j*_). Additionally, the number of bacteria and phage transferred from a neighbor node *j* to a local node *g* is re-scaled to the local density, i.e.,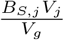.

Changes in the concentration of mucin lining the airways impact bacteria motility^29^ and phage diffusion^30,31^, thereby affecting their hopping rate throughout the bronchial network. In text S1 C-D, we explain in detail how we calculate the diffusion constants and the hopping rate of phage and bacteria across different mucin levels. Additionally, the diffusion values of phage and bacteria can be found in Table S2.

For phage infection, we use a model, *F* (*P*), that assumes spatial heterogeneities inside the mouse lungs might limit phage-bacteria encounters.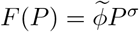, where 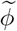 is the nonlinear phage adsorption rate and *σ <* 1 is the power-law exponent in the phage infection rate. This heterogeneous mixing model^13^ has been used previously to recapitulate phage-bacteria dynamics *in vivo*.

Text S1 includes additional details on the model, network structure, parameter estimation, and numerical simulation.

## III. RESULTS

### A. Spatiotemporal dynamics of phage therapy of acute pneumonia caused by *P. aeruginosa*

We begin by simulating phage therapy treatment of a *P. aeruginosa* infection in an immunocompetent host. The infection starts by inoculating the host with 10^6^ bacterial cells. For phage treatment, we add 10^7^ phage 2 hr after the beginning of the infection, which are parameters consistent with *in vivo* treatment of the MDR *P. aeruginosa* strain PAKlumi with the phage PAK P1^13^. The bacterial inoculum and the phage dose are uniformly distributed among network nodes, ensuring that each node has the same initial bacterial density of 1.11 × 10^6^ CFU/ml and phage density of 1.11 ×10^7^ PFU/ml, considering a network volume of ∼0.9 ml. We simulate baseline neutrophil levels by setting an initial immune density of 4.05 ×10^5^ neutrophils/ml^32^ in all network nodes.

By analyzing the dynamics that emerge from the phage therapy of a *P. aeruginosa* infection (Fig. 2), we observed that the phage-susceptible bacterial population (*B*_*S*_) initially grew and reached its peak density after 13 hr. The phage population (*P*) grew together with the *B*_*S*_ population during the first hours of the simulation. The host innate immune response (*I*) also increased in the presence of bacteria during the first 10 hr of the simulation. Once the phage population reached high-density levels, i.e., ∼10^11^ PFU/ml, we observed a reduction of the *B*_*S*_ population across the network. The bacterial elimination rate accelerated as the phage reduced the *B*_*S*_ population density to a level that facilitated the innate immune response to control the infection. Although phage-resistant mutants (*B*_*R*_) emerged and increased primarily in the top nodes of the network, the *B*_*R*_ population was maintained at levels that eased their control by the host innate immune response. The combined effects of phage and neutrophils led to the clearance of the infection in the network after 32 hr (Fig. 2), similar to observed clearance times of 24-48 hr in phage therapeutic treatment of *P. aeruginosa* in immunocompetent mice^13^. Note despite similar population dynamics between nodes, the infection clearance did not occur simultaneously in all the nodes. The infection resolved first in the bottom (∼28 hr) and later in the top nodes (∼33 hr).

**FIG. 2:**
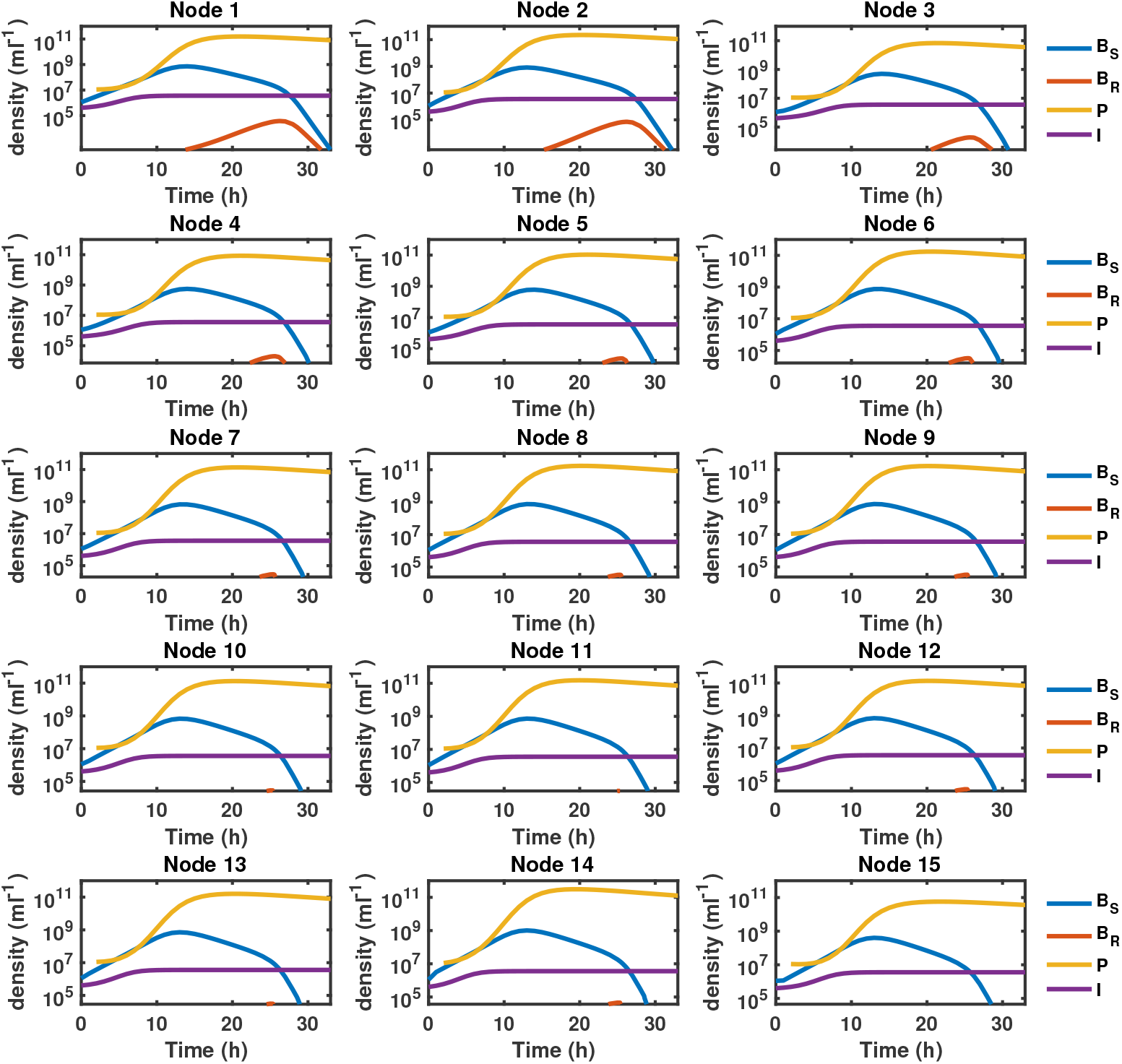
Spatiotemporal dynamics of phage therapy of a *P. aeruginosa* lung infection. We show the population dynamics at the node level of phage (solid yellow line), phage-susceptible bacteria (solid blue line), phage-resistant bacteria (solid orange line), and the host innate immune response (purple solid line). We infect an immunocompetent host with 10^6^ *P. aeruginosa* cells. Phage therapy (10^7^ PFU) is administered 2 hr after the bacterial inoculation. We uniformly distribute the phage dose and the bacterial inoculum in the network such that each node had the same initial bacterial density (1.11 × 10^6^ CFU/ml) and phage density (1.11 × 10^7^ PFU/ml). When the host is immunocompetent, we set the initial immune density to *I*_0_ = 4.05 × 10^5^ cells/ml in all the network nodes. The simulation runs for 33 hr. Here, Node 1 = Generation 1 = trachea, and Node 15 = Generation 15 = terminal airway.

### B. Joint action of phage and host innate immunity in bacterial infections

Next, we study how different treatment scenarios impact the bacterial infection spatiotemporal dynamics across the metapopulation network. We test four treatment scenarios that result from the presence (+) or absence (−) of both phage and innate immunity. As before, we inoculate a total of 10^6^ bacterial cells that are homogeneously distributed in the network. When phage therapy is applied, we inoculate the host with 10^7^ phage (MOI ≈10) 2 hr after the beginning of the infection. If the host is immunocompetent, we set an initial immune density of 4.05 × 10^5^ neutrophils/ml for all network nodes, totaling 3.64 ×10^5^ neutrophils in the lungs^32^. When the host is immunodeficient, we set the immune density to *I* = 0 neutrophils/ml in the network nodes. The bacterial inoculum and the phage dose are uniformly distributed in the network such that each node has the same initial bacterial density (1.11 × 10^6^ CFU/ml) and phage density (1.11 × 10^7^ PFU/ml), as in the prior section.

When an immunodeficient host was not treated with phage, bacteria grew unimpeded (Fig. S2a), reaching carrying capacity levels (1.5 × 10^9^ CFU/ml) after 10 hr in all the nodes (Fig. 3a). On the other hand, an active innate immune response delayed the initial growth of bacteria due to neutrophils killing the bacteria (Fig. 3b). However, in the absence of another antimicrobial effect, bacteria continued to grow and overwhelmed the innate immune response, rendering it insufficient for controlling the infection (Fig. S2b). The phage treatment of *P. aeruginosa* infection in an immunodeficient host led to eliminating the *B*_*S*_ population after 50 hr in most network nodes. However, it took about 90 hr to eliminate the *B*_*S*_ population in the top nodes (Fig. S2c). During this time, phage-resistant mutants (*B*_*R*_) emerged and spread throughout the network, causing the infection to persist (Fig. 3c). Finally, the model predicted therapeutic success when the host innate immunity complemented phage therapy. The predictions indicated eliminating bacteria from the network after 32 hr (Fig. 3d). The elimination of the infection followed a spatial pattern that started at the bottom and continued to the top network nodes (Fig. 3e). The observed pattern is compatible with a situation where phage help decrease the bacterial density to the point where the innate immune system is able to control and drive bacteria to extinction; refer to Text S1 E for additional analysis on the spatial structure in pathogen clearance. Next, we sought to characterize how changes in mucin levels—known for shaping phage and bacteria propagation rates^29–31^—impact the observed clearance pattern. The spatial clearance pattern was recapitulated for mucin levels varying from 1% to 4% (Fig. 3f). This suggests infection clearance is robust to changes in conditions that affect phage and bacteria propagation rates when the host is fully immunocompetent. Overall, modeling outcomes show that synergistic interactions between phage and neutrophils may lead to the resolution of the infection in spatially structured environments.

**FIG. 3:**
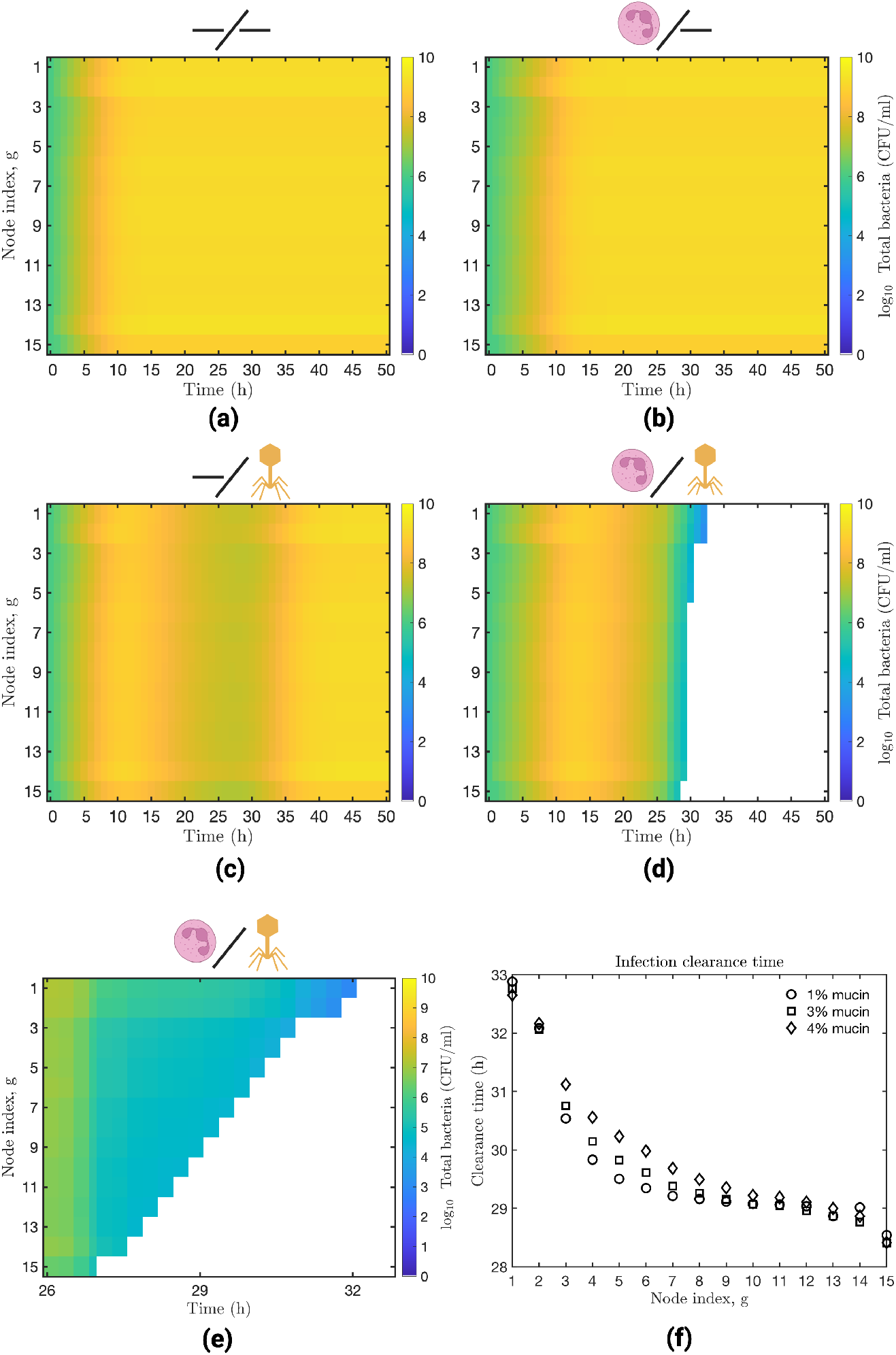
Bacterial dynamics under different phage and innate immune treatments. We simulate four treatment scenarios that result from the presence or absence (−) of both phage and the innate immune response. We show the bacterial dynamics across the metapopulation network when the host is immunodeficient untreated (a) or phage-treated (c). Similarly, we show the bacterial dynamics when the host is immunocompetent untreated (b) or phage-treated (d). The heatmaps depict the progression of the bacterial infection across the network; each row represents a network node, *g*, while the columns indicate the simulation time (hr). The node color represents the bacterial density at a given time. The yellow regions represent high bacterial density, and the white areas represent infection clearance. When the host is immunocompetent and phage-treated, we zoom in and show the infection clearance pattern (e). We also test the effects of varying the mucin level (1-4%) on the infection clearance time (f). In all simulations, we inoculate a host with 10^6^ bacterial cells. If phage therapy is used, we administer 10^7^ phage (PFU) 2 hr after the beginning of the infection. We uniformly distribute the phage dose and the bacterial inoculum such that each node has the same initial bacterial density (1.11 × 10^6^ CFU/ml) and phage density (1.11 ×10^7^ PFU/ml). When the host is immunocompetent, we set an initial immune density of 4.05 × 10^5^ cells/ml in all the nodes. If the host is immunodeficient, we set the immune density to *I*_0_ = 0 cells/ml. A 2.5% mucin concentration was used for scenarios (a) to (e). All the simulations ran for 50 hr. On the heatmaps, row 1 represents Generation 1 and the top of the lungs, while row 15 represents Generation 15 and the bottom of the lungs.

### C. Varying the allocation of bacterial inoculum and phage dose in the network does not affect the therapeutic outcome

Thus far, we have explored the impacts of phage therapy on a *P. aeruginosa* infection by uniformly distributing the bacterial inoculum and the phage dose in the network. However, the actual distribution of phage and bacteria in the bronchial tree after intranasal inoculation is not necessarily well-constrained. Hence, we try different forms of allocating the bacteria and phage inocula in the metapopulation network. For example, (1) we uniformly distribute the bacterial inoculum such that every node has the same initial density (uniform distribution), (2) we distribute the inoculum among the first three nodes of the network (top distribution), (3) we distribute the inoculum among the last twelve nodes of the network (bottom distribution), and (4) we only inoculate the first node of the network (i.e., intratracheal instillation). We use the same distribution forms as the bacterial inoculum for the phage dose. In total, we test 16 ways of distributing the bacterial inoculum and the phage dose in the network (Fig. 4a).

**FIG. 4:**
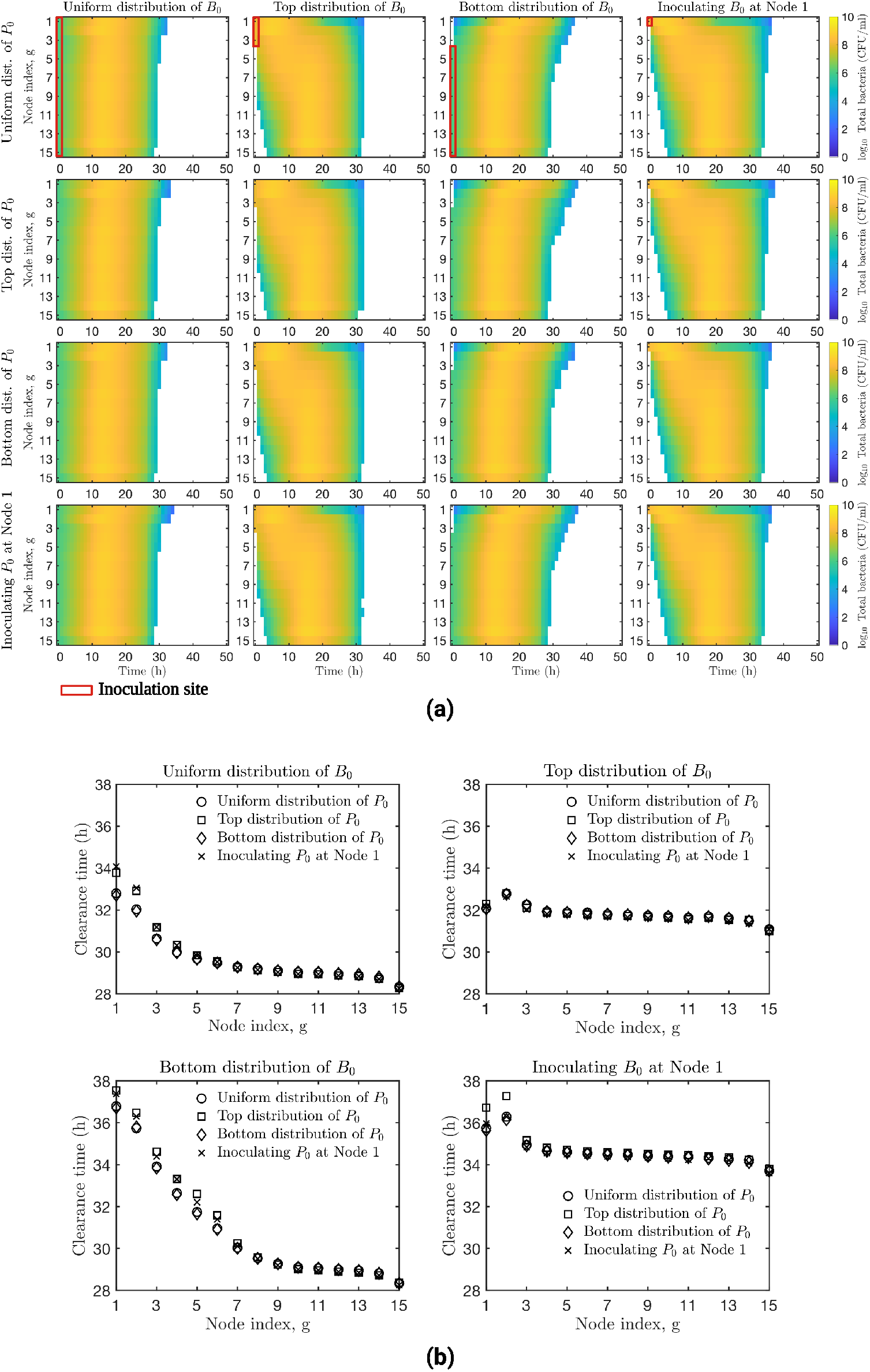
Infection dynamics as a result of varying the distribution of phage dose and bacterial inoculum in the network. We evaluate different forms of allocating the bacterial inoculum (*B*_0_) and the phage dose (*P*_0_) among network nodes (a). For example, 1) we uniformly distribute the phage dose or the bacterial inoculum among the network nodes, 2) we distribute the phage dose or the bacterial inoculum between the first three nodes of the network (Top distribution), 3) we distribute the phage dose or the bacterial inoculum among the last 12 nodes of the network (Bottom distribution), or 4) we inoculate the first node of the network with phage or bacteria. We use a heatmap to represent paired distributions of phage dose and bacterial inoculum. We show how bacterial infection progresses per network node (g). In the heatmap, each row represents a network node, while the columns indicate the simulation time. The node color represents the bacterial density, the yellow regions represent high bacterial density, and the white areas represent infection elimination. Given a pair of bacterial inoculum and phage dose distributions, we calculate the infection clearance time at the node level (b). For (a) and (b), we infect an immunocompetent host with 10^6^ bacterial cells. We administer 10^7^ phage (PFU) 2 hr after the bacterial infection. We set an initial immune density of *I*_0_ = 4.05 ×10^5^ cells/ml in all the nodes. The simulation ran for 50 hr. On the heatmaps, row 1 represents Generation 1 and the top of the lungs, while row 15 represents Generation 15 and the bottom of the lungs.

When the bacterial inoculum is uniformly distributed, the model predicted the elimination of bacteria 32 to ∼34 hr after the beginning of the infection, regardless of how phage dose was allocated in the network (Fig. 4a-1st left-column). When the first three nodes were inoculated with bacteria, it took ∼4 hr for bacteria to colonize the bottom of the network and around ∼32 hr to eliminate the infection from the network regardless of phage dose distribution (Fig. 4a-2nd column). The delay in colonizing bottom nodes led to delayed elimination of the infection in those nodes; e.g., compare clearance times of top vs. uniform distribution of *B*_0_ in Fig. 4b. When the bacterial inoculum was distributed among the last twelve nodes of the network, it took ~37 hr to clear the infection from the network. The colonization of bacteria in the top nodes was delayed due to the initial distribution of the inoculum (Fig. 4a-3rd column). In this scenario, there was a longer infection clearance time difference between the top (clearance occurred at 37 hr) and the bottom (clearance occurred at 28 hr) of the network (Fig. 4b-Bottom distribution of *B*_0_). Finally, the inoculation of bacteria in node one resulted in the colonization of bottom nodes after ~6 hr and elimination of the infection after 36 hr (Fig. 4a last right-column). There was a delay in removing the bacterial infection from the bottom nodes, similar to the previous scenario where the inoculum was distributed only among the top nodes. We observed different bacterial colonization patterns depending on how the bacterial inoculum was initially distributed in the network. Varying the allocation of the phage dose in the network did not significantly impact bacterial colonization or infection clearance patterns. As in previous simulations (Fig. 2 and Fig. 3e), we noticed a consistent spatial pattern of infection resolution across all phage and bacteria inocula distributions, where the infection resolved faster at the bottom than at the top of the network (see Text S1 E for details of the spatial pattern of infection elimination).

### D. Sufficient neutrophil levels and phage adsorption rates are required for effective clearance of *P. aeruginosa* lung infection

As part of our model analysis, we study how intermediate phage efficacy and host innate immunity states impact metapopulation model outcomes, comparing the results with those from the well-mixed case. To model intermediate immune states, we vary the percentage of neutrophil availability in the lungs from 1% to 100%, where 100% availability corresponds to ∼3.24 ×10^6^ lung neutrophils in immunocompetent mice^32^. To explore intermediate phage efficacy, we vary phage adsorption rate across a range from 10^−9^ to 10^−6^ (ml/PFU)^*σ*^*h*^−1^. We simulate the phage treatment of a *P. aeruginosa* infection by inoculating a host with 10^6^ bacterial cells and introducing 10^7^ phage 2 hr after the bacterial infection. To assess the robustness of model predictions, we randomize the initial conditions and try 84 different ways of allocating the bacterial inoculum and the phage dose in the network. Then, we calculate the probability of clearing the infection by simulating the different initial conditions, given a specific immune state and phage adsorption rate.

When neutrophil availability is limited (e.g., 10-35%), the metapopulation model predicted between 25% to 42% the chances of therapeutic success for phage adsorption rates 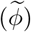 ranging between ∼ 2.5 ×10^−7^ to 10^−6^ (ml/PFU)^*σ*^*h*^−1^ (Fig. 5). In contrast, the well-mixed model, lacking consideration of spatial structure, predicts infection persistence in the same parameter range. The spatial model predicts between 40% to 100% the probabilities of therapeutic success when 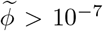 (ml/PFU)^*σ*^*h*^−1^ and for neutrophil availability *>*50%. In contrast, infection was always resolved in the well-mixed model within the specified parameter range (Fig. 5, region limited by solid white line). For example, if we compare the metapopulation model predictions for a 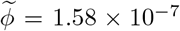 (ml/PFU)^*σ*^*h*^−1^, neutrophil availability should be *>*85% to have increased chances of therapeutic success (i.e., *>*60%), while for the same 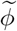 value, infection always clears in the well-mixed model when neutrophil availability is *>*55%. These findings highlight the role of spatial structure in shaping phage therapy outcomes and suggest that higher levels of host innate immunity might be required to successfully clear a *P. aeruginosa* infection when spatial constraints are considered.

**FIG. 5:**
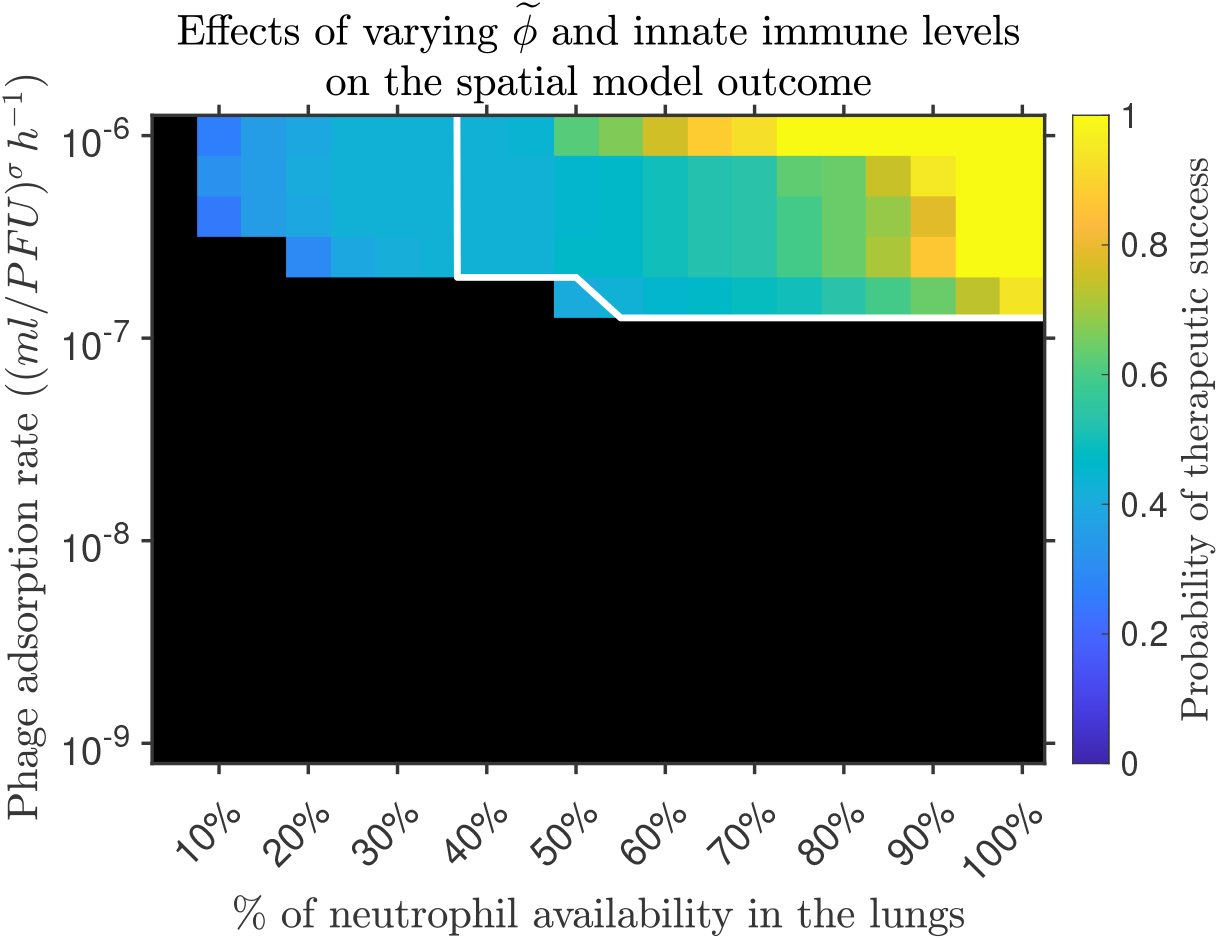
Probability of therapeutic success given intermediate phage efficacy and host innate immune levels. To explore intermediate innate immune responses, we vary the percentage of neutrophils available in the lungs from 1 to 100%. We also test intermediate phage efficacy by varying the phage adsorption rate within the range 10^−9^ to 10^−6^ (ml/PFU)^*σ*^*h*^−1^. We test the robustness of the model by randomizing the initial conditions and trying 84 different ways of distributing phage dose and bacterial inoculum across the network. Then, we calculate the probability of clearing the infection by simulating the 84 initial conditions under specific phage adsorption rates and immune response levels. The heatmap shows the probability of clearing the infection for the specified phage adsorption rates and immune response levels. The colored regions represent a *p >* 0 of clearing the infection, while black regions represent failure to clear the infection, i.e., a *p* = 0 of therapeutic success. The white solid line contours the region of infection clearance predicted by the well-mixed model. To simulate the phage treatment of a *P. aeruginosa* infection, we inoculate a host with 10^6^ bacterial cells and introduce 10^7^ phage 2 hr after the bacterial infection. The simulation runs for 250 hr. A 100% neutrophil availability represents a total of 3.24 ×10^6^ lung neutrophils^32^. In this simulation, we use a 2.5% mucin level.

Within the metapopulation model outcomes, we noticed a trend where the higher the neutrophil availability the wider the range of 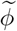 values for which the model predicts increased chances of infection clearance. This suggests that despite spatial structure effects that may affect phage-bacteria encounters and the phage adsorption rate, having a robust immune response may facilitate the control of the *P. aeruginosa* infection. On the other hand, low phage efficacy, e.g., 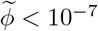 (ml/PFU)^*σ*^*h*^−1^, may render phage therapy inefficient even when the host is fully immunocompetent, according to predictions from both the metapopulation and well-mixed models (Fig. 5, black region).

Note that we also explored how variations in mucin and innate immune levels impact phage therapy outcomes. The exploratory analysis confirms the synergistic elimination of the infection by phage and neutrophils given sufficiently active immune states and depicts a weak effect of mucin level variation (Fig. S4/Text S1 F).

### E. Analysis of *in vivo P. aeruginosa* infection data confirms a bottom-to-top infection clearance pattern

Our finding of a systematic bottom-to-top transition pattern supported by multiple simulations prompted us to examine spatial patterns of clearance within *in vivo* imaging data from phage treatment of *P. aeruginosa* infected mice. We analyzed the bioluminescence infection signal from images (Fig. 6) generated in a previous phage therapy study^13^. The dataset included images of 13 mice, comprising both WT (N = 4) and lymphocyte-deficient *Rag*2^−*/*−^*Il*2*rg*^−*/*−^ (N = 9) mice groups. Following *P. aeruginosa* infection (10^7^ CFU), mice were treated with phages (10^8^ PFU) 2 hr later. We split individual mouse images into two compartments (dashed white line in Fig. 6). The top compartment encompassed the nose and throat, while the bottom compartment included the lungs. To quantify infection intensity within each compartment, we computed the total intensity signal by summing the pixel intensity values of all pixels within a compartment (Fig. 7a). This total intensity served as a proxy for the bacterial infection status inside the mouse, where higher values indicated higher bacterial density. To determine infection signal clearance in each compartment, we established an intensity threshold of 3, below which we considered the total intensity signal cleared. For additional details of the image analysis methodology, refer to Text S1 G.

**FIG. 6:**
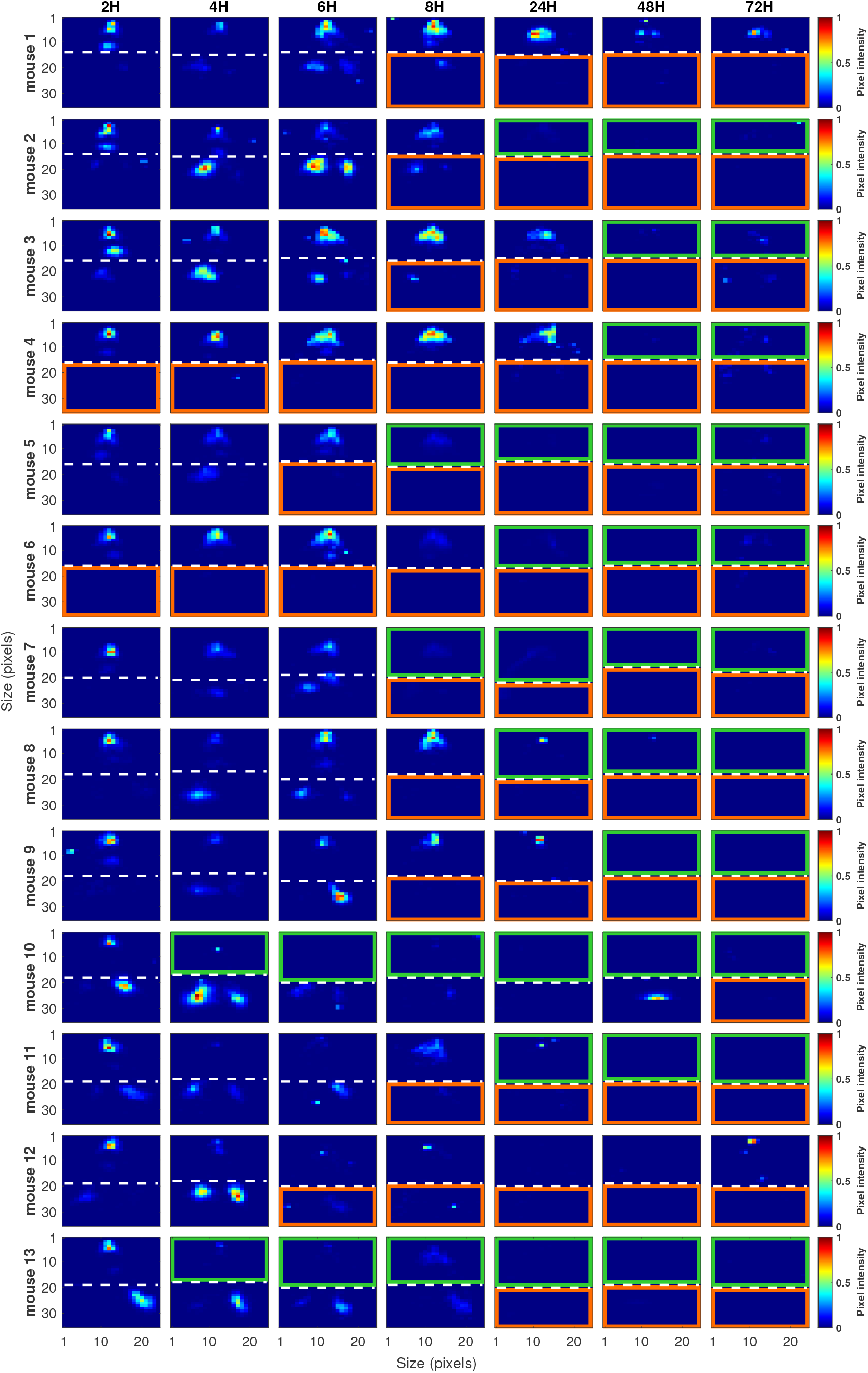
*In vivo P. aeruginosa* murine pneumonia data. We show mice images depicting the evolution of a *P. aeruginosa* infection *in vivo* during 72 hr. We show data for two mice groups, WT (N = 4; mouse #1 to 4) and *Rag*2^−*/*−^*Il*2*rg*^−*/*−^ (N = 9; mouse #5 to 13). The intensity of the bioluminescence signal represents the intensity of the infection in different mouse regions. The pixel intensity value is our proxy for the bacterial density. A pixel intensity of 1 represents the highest bacterial density, while 0 represents the threshold of detection. The white dashed line separates the upper and lower compartments of the mouse respiratory system. The orange and green boxes highlight the approximate time when the total intensity signal drops below the intensity threshold in the lower and upper compartments, respectively. Mice were inoculated with 10^7^ *P. aeruginosa* cells, and 2 hr after the bacterial inoculation, mice were treated with phage (10^8^ PFU).

**FIG. 7:**
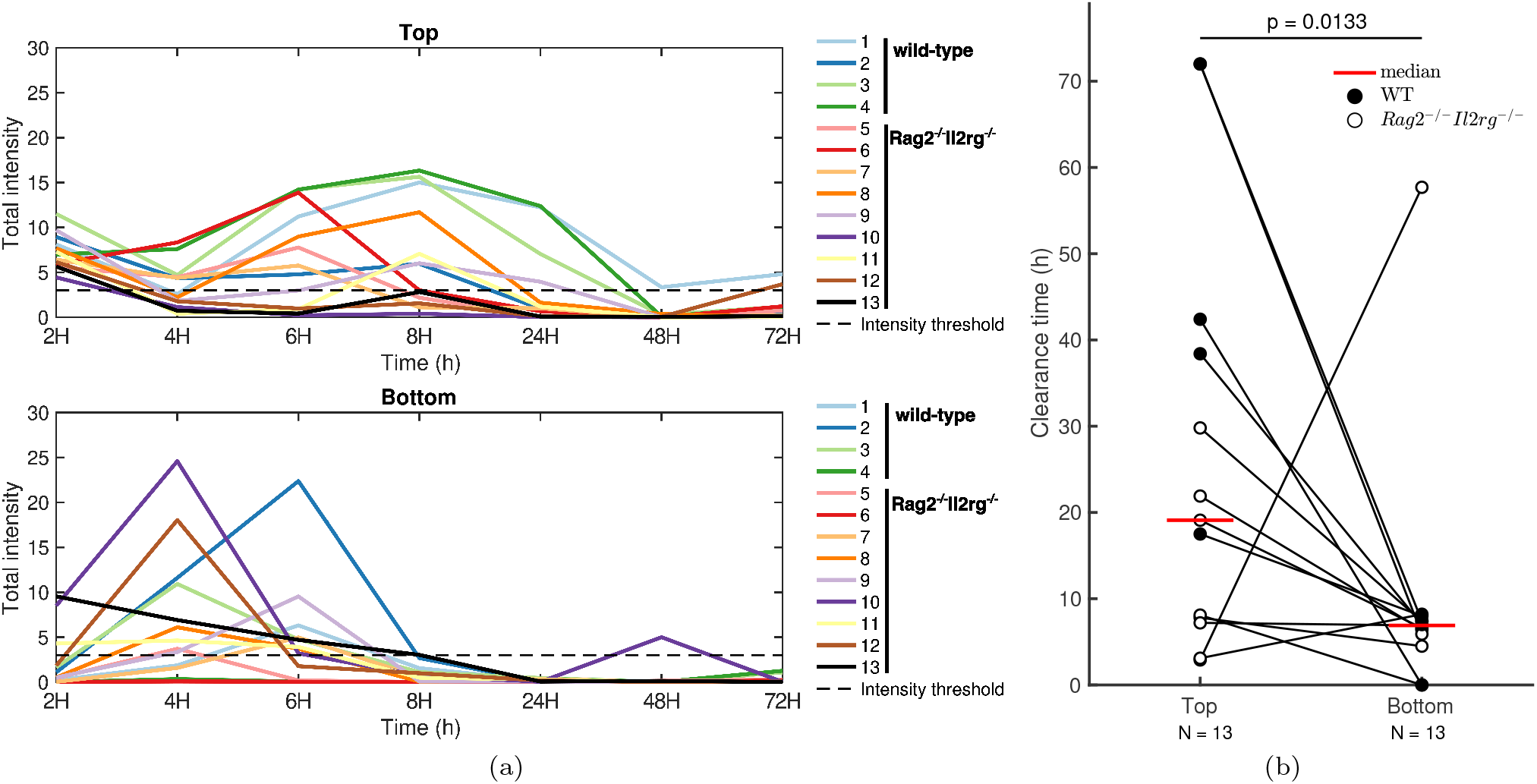
Time series of total intensity signal and the infection clearance analysis using *in vivo P. aeruginosa* murine pneumonia data. We show the time series of total intensity signals for the upper and lower compartments of 13 mice (a). The total intensity of one compartment is calculated by adding the pixel intensity values from all pixels making up a compartment. The black dashed line represents the intensity threshold below which the total intensity signal clears. We calculate the time to infection resolution for the upper and lower compartments (b). We use data from 13 mice, including WT (N = 4) and *Rag*2^−*/*−^*Il*2*rg*^−*/*−^ (N = 9) mice groups. We used the one-sided Wilcoxon signed rank test to compare the infection clearance time difference between the upper and lower compartments (b).

Using the time series of imaged mice, we tracked the progression of the infection in the two compartments (Fig. 7a). We observed a more rapid decrease in intensity signal below the threshold in the bottom compartment (∼6-8 hr after the onset of infection) compared to the top compartment (∼8-48 hr after the onset of infection). The observation implies a faster clearance of infection in the lower respiratory tract than in the upper respiratory tract of mice. Upon calculating the infection clearance time for both compartments, we found that the intensity signal cleared faster in the bottom compartment than in the top compartment. Specifically, the median time to infection clearance was 7 hr for the bottom compartment and 19 hr for the top compartment, yielding a statistically significant difference with a p-value of 0.0133 (Fig. 7b). These spatial patterns of infection resolution align with the qualitative predictions made by the metapopulation model, particularly under conditions involving active innate immunity and phage therapy.

## IV. DISCUSSION

In this study, we developed a metapopulation model of phage therapy of a *P. aeruginosa* lung infection. We modeled the ecological interactions between phage, two *P. aeruginosa* strains (phage-susceptible and phage-resistant), and the host innate immune response at the airway level. The dynamics arising from the phage therapy of a *P. aeruginosa* infection in an immunocompetent host indicates that phage therapy successfully eliminates bacterial infections when the host innate immune responses complement phage treatment. The therapeutic outcome was robust to heterogeneous distributions of phage dose and bacterial inoculum in the network and to variations in mucin levels. However, limited neutrophil availability and a low phage adsorption rate (e.g., *<* 10^−7^ (ml/PFU)^*σ*^*h*^−1^) may negatively impact phage treatment efficacy. Moreover, the metapopulation model required higher innate immune response levels to increase the likelihood of therapeutic success, compared to a well-mixed model that achieved infection resolution at lower innate immune levels, highlighting the role of spatial structure in shaping phage therapy outcomes. Throughout the simulations, we note a spatial pattern of infection clearance, wherein infection resolved faster in the bottom compared to the top nodes. The spatial clearance pattern results in part from dynamics crossing a critical threshold via a volumetric effect associated with the smaller total number of bacterial cells in bottom vs. top nodes. Finally, by analyzing *in vivo* data of mice infected with *P. aeruginosa* and treated with phage, we identified spatial transitions in pathogen clearance that align with the metapopulation model outcomes.

The therapeutic outcome recapitulated in our model is consistent with *in vivo* phage therapy results, where phage efficiently controlled respiratory bacterial infections in both humans^33–35^ and mice^13,36–38^. Notably, we found that the therapeutic outcome was robust to different distributions of the phage dose across the bronchial network, including heterogeneous and homogeneous dose distributions. This is consistent with outcomes from phage delivery strategies using phage-loaded microparticles to distribute phages throughout the lung, effectively reducing infections caused by *P. aeruginosa*^39^ and *S. aureus*^40^. Furthermore, Delattre et al. developed a computational model that recapitulated phage-bacteria interactions *in vivo* and found that varying the route of administration of phage dose, i.e., intratracheally or intravenously, did not impact the therapeutic outcome with both courses producing similar effects^36^. While variations in bacterial inoculum allocation impacted colonization patterns within the network, they did not disrupt the therapeutic outcome. These collective findings suggest that therapeutic success might be robust to initial phage and bacterial distributions in the lungs, though quantitative differences can rise with mismatched distributions.

Immunophage synergy does have its limits, whether *in vivo*^13^, in mean field models^19^, or as shown here, in metapopulation models. For example, a weak innate immune response may negatively impact phage treatment efficacy. Our model predicted a minimum of 45% of neutrophil availability to start clearing bacteria from the lungs, although higher innate immunity levels are needed to increase the likelihood of therapeutic success. This is consistent with *in vivo* experiments showing that phage therapy failed to control a *P. aeruginosa* infection when mice had an impaired innate immune system, e.g., when mice were either MyD88-deficient^13^ or neutropenic^13,14^. On the other hand, phage therapy successfully eliminated the infection when mice were immunocompetent^13,14,37^. Future phage therapy studies and clinical trials should examine the importance of the host immune status in determining treatment efficacy.

The metapopulation network model has some caveats. For example, the network structure of the model is based on a symmetrical bronchial tree, while the mouse bronchial tree is asymmetric^41^. Hence, future model extensions could consider modeling asymmetric branching structures and their effects on phage and bacteria propagation across the bronchial tree. By doing so, we can evaluate if tree asymmetries facilitate spatial refuges for bacteria to protect themselves from phage attacks. A critical defense mechanism of the respiratory system is mucociliary clearance^42^ (MCC), where motile cilia of the airway epithelium transport a mucus layer out of the lungs, carrying particles trapped in the mucus layer. Currently, our model does not address the mechanism by which bacteria first infect the lungs, the mucosal layer’s involvement in this process, or the potential translocation of bacteria between different body sites^43^. In the future, we could extend the model to include MCC effects to assess how MCC impacts bacterial colonization, infection clearance patterns, and neutrophil transport within the airways. Our model does not account for the interactions between pathogenic *P. aeruginosa* and the lung microbiome^44^. Furthermore, we do not consider the effects of other innate effector cells, such as macrophages, the predominant immune cells in the mice lungs before infection begins^32^. Alveolar macrophage’s role in the activation of neutrophils, phagocytosis, and the possible interactions these cells have with phage^45,46^ should be considered for future model extensions.

Phage therapy has emerged as an alternative to chemical antibiotics to treat infections by MDR bacteria. However, spatial structure effects of *in vivo* environments where phage and bacteria interact have not been fully addressed yet. We used a metapopulation network model to study the impact of spatial structure effects on phage therapy *in vivo*. Synergistic interactions between phage and neutrophils lead to bacteria elimination within the lung environment. However, higher innate immune levels could be required to increase the likelihood of therapeutic success in a spatially structured environment compared to a well-mixed system. Our modeling outcomes further demonstrated the importance of the host immune status and phage life history traits, such as phage adsorption rate, in shaping the therapeutic outcome. The emergence of lung infection spatiotemporal clearance patterns also suggests that extending *in vitro* phage therapy models may help to guide therapeutic treatment development in other *in vivo* contexts.

## V. ACKNOWLEDGMENTS

The work was supported by the National Institutes of Health (R01 AI146592 to JSW, LD). JSW was supported, in part, by the Chaires Blaise Pascal program of the Île-de-France region. RRG was supported, in part, by the National Council of Humanities, Science, and Technology of Mexico. We thank Jeremy Seurat for his insights on the statistical analysis part of this manuscript. We thank Stephen Beckett for his feedback on the metapopulation network structure and the manuscript.

## Supplementary Text S1: Model details

We determine the number of generations (*N*) that the metapopulation network must have, based on the volume of the network (*V*_*network*_) and the total lung capacity (TLC) of mice^27^, which is approximately 1 ml. To do so, we consider the volume of individual airways (*V*_*airway,g*_) and the number of airways present at each generation, ∼ 2^*g*−1^. Then, we find *N* by approximating *V*_*network*_ to the TLC of the mouse.

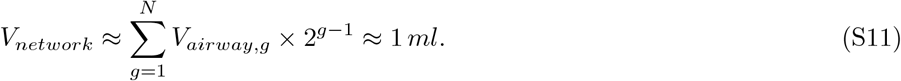

We find that with *N* = 15, we do not exceed the TLC of the mouse, and we obtain a network volume of ∼0.9 ml. We use 15 generations (nodes) for the metapopulation network model.

### A. Model simulation

Model equations (Eqns. 1-4) are numerically integrated using ODE45 in MATLAB R2020b. By doing so, we obtain the population dynamics of phage-susceptible and phage-resistant bacteria, phage, and the innate immune response. We assume bacteria become extinct in a node when the number of bacteria drops below the extinction threshold of 1 CFU. This applies to both *B*_*S*_ and *B*_*R*_. When bacterial counts drop below the extinction threshold in a node, we set the bacterial density to 0 CFU/ml.

### B. Parameter estimation

The parameter values used in the simulations of the metapopulation model are shown in Table S1. Most of the parameter estimation was carried out in previous work (see “Parameter Estimation” section in reference^13^). Airway anatomical information, including airway length and diameter, was obtained from^28^. Bacteria and phage diffusion constants can be found in Table S2.

**TABLE S1:** Parameter values of the metapopulation network model. Concentrations are calculated based on mice lung volume of 0.9 ml.

| Parameters of the metapopulation model | Value | Estimated from |
| --- | --- | --- |
| $r_P$, maximum growth rate of phage-susceptible bacteria | $0.75 \text{ h}^{-1}$ | *P. aeruginosa* murine pneumonia model<sup>47</sup> |
| $r_A$, maximum growth rate of phage-resistant bacteria | $0.675 \text{ h}^{-1}$ | 10% tradeoff between resistance against phage and growth rate<sup>48</sup> |
| $K_C$, carrying capacity of bacteria | $1.49 \times 10^9 \text{ CFU/ml}$ | Estimated from<sup>13</sup> infection |
| $\beta$, burst size of phage | 100 | Estimated from<sup>13</sup> |
| $\omega$, decay rate of phage | $0.07 \text{ h}^{-1}$ | Estimated from<sup>13</sup> |
| $\epsilon$, killing rate parameter of immune response | $5.47 \times 10^{-7} \text{ ml/(h cell)}$ | Set such that $\epsilon K_I$ gives the maximum granulocyte killing rate<sup>47</sup> |
| $\alpha$, maximum growth rate of immune response | $0.97 \text{ h}^{-1}$ | Fitting of neutrophil recruitment data<sup>32</sup> |
| $K_I$, maximum capacity of immune response | $3.59 \times 10^6 \text{ cell/ml}$ | Fitting of neutrophil recruitment data<sup>32</sup> |
| $K_D$, bacteria concentration at which immune response is half as effective | $6.14 \times 10^6 \text{ CFU/ml}$ | Corresponds to a lethal dose of about $5.5 \times 10^6 \text{ CFU/lungs}$ |
| $K_N$, bacteria concentration when the immune response growth rate is half its maximum | $10^7 \text{ CFU/ml}$ | In vitro data of TLR5 response to PAK strain<sup>49</sup> |
| $B_0$, initial bacterial density (when inoculum is uniformly distributed among network nodes) | $1.1106 \times 10^6 \text{ CFU/ml}$ | Total inoculum of $10^6 \text{ CFU}$ |
| $P_0$, initial phage dose (when dose is uniformly distributed among network nodes) | $1.1106 \times 10^7 \text{ PFU/ml}$ | Total phage dose of $10^7 \text{ PFU}$ |
| $I_0$, initial immune response | $4.048 \times 10^5 \text{ cell/ml}$ | Fitting of neutrophil recruitment data<sup>32</sup> |
| $I_0$, initial immune response (neutropenic mice) | $0 \text{ cell/ml}$ | Assuming no primary innate immunity |
| $N_{Lung}$, total number of neutrophils in the lungs | $3.24 \times 10^6 \text{ cells}$ | Fitting of neutrophil recruitment data<sup>32</sup> |
| $\mu_1$, probability of emergence of phage-resistant mutants per cellular division | $2.85 \times 10^{-8}$ | Estimated from experimental measurements<sup>50</sup> |
| $\mu_2$, probability of emergence of reversible mutant (i.e., from phage-resistant to phage-susceptible) per cellular division | $2.85 \times 10^{-8}$ | Approximated to the estimates from<sup>50</sup> |
| **Heterogeneous mixing (HM) model parameters** |  |  |
| $\phi$, nonlinear phage adsorption rate | $1.686 \times 10^{-7} \text{ (ml/PFU)}^\sigma \text{ h}^{-1}$ | Estimated from<sup>13</sup> |
| $\sigma$, power law exponent in phage infection rate | 0.6 | Estimated from<sup>13</sup> |
| **Anatomical information of airways** |  |  |
| $V_g$, airway volume | $4.8 \times 10^{-3} - 2.2 \times 10^{-5} \text{ ml}$ | Calculated as the volume of a cylinder ($\pi r^2 h$), using anatomical information of mice airways from<sup>28</sup> |
| $l_g$, airway length | 0.76 - 0.061 cm | Anatomical information of mice airways from<sup>28</sup> |
| airway diameter | 0.089 - 0.02 cm | Anatomical information of mice airways from<sup>28</sup> |

### C. Calculating bacteria and phage diffusion constants as a function of mucin level

The hopping rate is calculated as the reciprocal of the time it would take for bacteria and phage to cross half of the airway via diffusion, *D*. Hence, we need to determine the diffusion constants of phage and bacteria to calculate their hopping rate. The evidence shows that the diffusion constant of phage^30,31^ and the speed of bacteria^29^ are shaped by the concentration of the mucus lining the airways. For example, highly concentrated mucus can reduce bacteria motility and slow phage diffusion. Our model focuses on mucin levels that are physiologically relevant in acute lung infections (e.g., 0-4% mucin concentration). Below, we explain how we calculate the diffusion coefficients of phage and bacteria across different mucin levels.

We collected speed values of *P. aeruginosa* for two different mucin levels^29^ (2.5% and 8%). We use the speed values to calculate the diffusion coefficient of bacteria using the run-and-tumble model as a proxy,

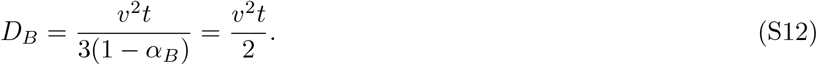

Where *v* is the bacterial speed in *µm/s, t* = 1 *s*, is the duration of the bacterial runs, and *θ* ∼60 degrees, is the average change in direction between runs, such that *α*_*B*_ = ⟨cos(*> θ*) ⟩ ≈1*/*3.

To find additional bacteria speed values within our target range of mucin concentrations (0-4%), we fit a line between the data points we collected (Fig. S1a). By calculating the parameters of the linear function, we can find the speed of bacteria for different mucin levels relevant to our biological system. Then, we calculate the bacteria diffusion coefficient from the speed value using the run-and-tumble model (Eqn. S12).

**TABLE S2:**
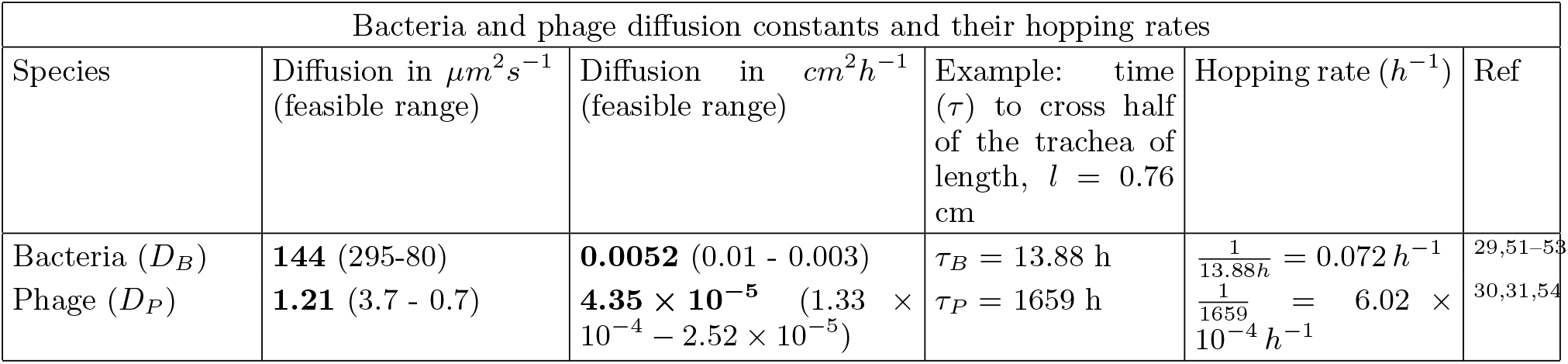
Examples of phage and bacteria diffusion coefficients under normal mucus concentration (2.5%). In parentheses, we show a feasible range of diffusion values when mucin levels vary from 0 to 4%.

We calculate the phage diffusion constant across different mucin levels by using the Stokes-Einstein equation (Eqn. S13). To do so, we use the predicted values of mucus viscosity (*η*_*M*_) across different mucin levels using the formula, *η*_*M*_ = *a*· [*mucin*]^*b*^ + *η*_*W*_, obtained from^31^. In that study, they fitted the mucus viscosity to empirical data using two power functions for two different regions of mucin levels, low [0-1% mucin] and high [1-4% mucin] concentrations. The parameter *a* has values of 0.326 and 0.323 in the low and high mucin regions, respectively. The exponent *b* has values of 0.65 and 1.59 for the low and high mucin regions, respectively. Parameters *a* and *b* can also be found in Tables 3 and 4 of the study^31^.

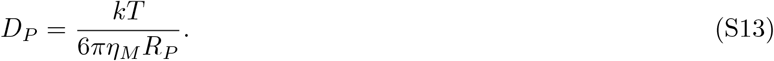

For the phage diffusion constant calculations, thermal energy was assumed at 37°C such that *kT* = 4.28 *pN* · *nm* and water viscosity was *η*_*W*_ = 0.69 *mPa* · *s*. The phage radius (*R*_*P*_) was calculated to be 90 *nm*, half the length of a typical T4 phage^31^. Using the Stokes-Einstein equation (Eqn. S13) and the predicted values of mucus viscosity across different mucin levels, we can calculate the phage diffusion constant for different mucin levels (Fig. S1b).

Using the relationship between mucin levels and phage and bacteria diffusion coefficients, we can explore how lung physiological conditions impact mucus concentration and affect the infection dynamics. For example, variations in the concentration of mucin could impact the time it takes for bacteria to colonize different nodes at various depths of the bronchial tree, phage dispersion times, the establishment of phage resistance, and the clearance of the infection by the combined effects of phage and neutrophils at different depths of the bronchial tree.

### D. Calculating the hopping rate of bacteria and phage

We calculate the hopping rate of bacteria and phage using the airway length information^28^ and the diffusion constants of phage and bacteria (Table S2). The hopping rate is calculated as the reciprocal of the time, *τ*, it would take for each species to cross half of the airway, *l/*2, via diffusion, *D*. We leverage the relationship between the mean squared displacement (MSD) and 1D Brownian motion, ⟨(*x* − *x*_0_)^2^⟩ = 2*Dt*, to calculate *τ*_*B*_ and *τ*_*P*_.

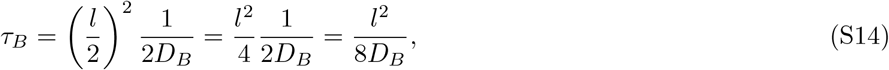

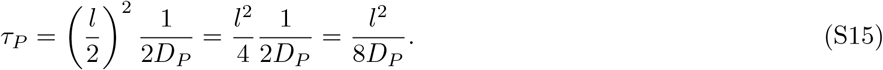

Then, we take the reciprocal of *τ*_*B*_ and *τ*_*P*_ to calculate the hopping rates of bacteria and phage, respectively. The hopping rate determines how fast bacteria and phage leave a local airway and hop to a neighboring airway. We use the hopping rates to calculate the influx and outflux terms of equations 1-3.

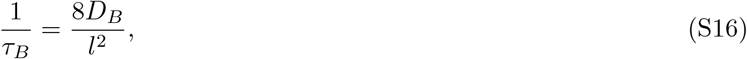

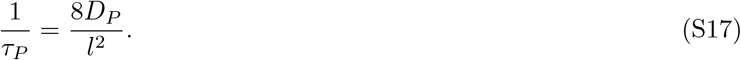

### E. Spatial pattern of infection elimination, innate immunity effects

The synergistic clearance of *P. aeruginosa* infection by phage and host innate immunity yielded a recurring feature: a spatial pattern of clearance of pathogens from bottom-to-top of the bronchial network. This pattern is compatible with a situation in which phage initially help decrease bacterial density to the point where the immune system alone can control and drive bacteria to extinction. Once bacteria fall below a critical level (*K*_*D*_), the immune system removes bacteria at a fixed rate until complete elimination within each node. We hypothesize that the observed difference in clearance times between the bottom and top nodes is influenced, in part, by node volume differences. Our model associates the bacterial extinction threshold with the number of bacterial cells crossing the 1 CFU threshold at the node level, suggesting that the spatial pattern in pathogen clearance could be a result of dynamics crossing a critical threshold via a volumetric effect associated with the smaller total number of bacterial cells in bottom vs. top nodes. If this were the case, the clearance time at the node level could be predicted using the immune killing rate (*εK*_*I*_), providing insights into the spatial variations observed in the metapopulation model simulations.

When calculating clearance time, we make several assumptions, including that *B*_*S*_ is the predominant bacterial type throughout the simulation, that bacterial density levels are below a critical threshold (*B*_*S*_ *<< K*_*D*_) at final simulation times, and that host innate immunity reached its maximum level (*K*_*I*_) by the end of the simulation. These conditions have been met in prior simulations (Fig. 2) depicting phage therapy for a *P. aeruginosa* infection in an immunocompetent host. Moreover, it is important to note that low bacterial density levels (*B*_*S*_ *<< K*_*C*_) enable positive exponential growth (*r*_*P*_ *B*_*S*_), and such growth should be considered when calculating the infection clearance time.

To determine the time to bacterial extinction, we solve the 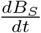 equation describing how *B*_*S*_ population changes over time due to the innate immune response and exponential growth. After solving the equation, we isolate *t* to determine the clearance time.

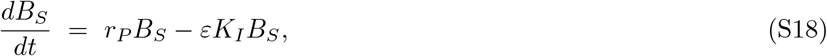

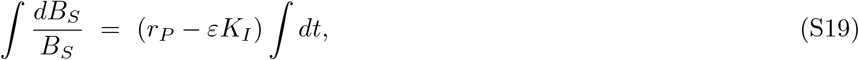

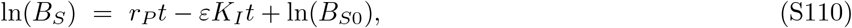

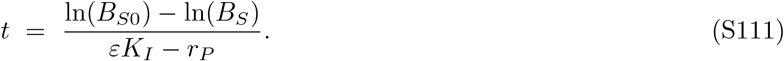

Bacteria become extinct in node *i* once their levels fall below the 1/*V*_*i*_ density threshold, where *V*_*i*_ is the volume of node *i*. So, the time to bacterial extinction, denoted as *t*_*i*_, for node *i* is

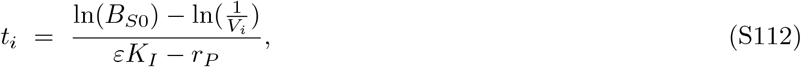

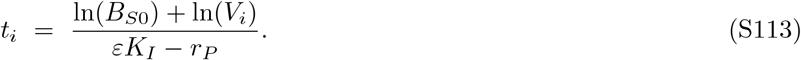

We note that time to infection resolution in node *i* is proportional to the natural logarithm of the node volume (*V*_*i*_) and scales with the inverse of the immune killing rate adjusted by the per capita bacterial growth rate (*r*_*P*_).

To further characterize the infection clearance time (ICT) difference between nodes, we calculate the ICT difference between node *i* and the last network node (which is the node with the smallest volume), i.e., *t*_*i*_ − *t*_*bottom*_. To do so, we use the clearance time calculated in Eqn. S113. The ICT difference between nodes due to the immune killing rate is,

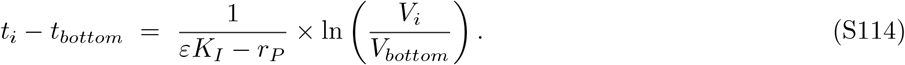

We observe that the clearance time difference is proportional to the natural logarithm of the volume ratio, *V*_*i*_*/V*_*bottom*_, where *V*_*bottom*_ is the volume of the last network node. We conclude that a larger volume difference between nodes corresponds to a longer clearance time difference. The largest volume difference in the network occurs between node one (trachea) and node fifteen (terminal airway), resulting in the longest ICT difference between any pair of nodes. Comparing the theoretical ICT difference to that obtained from simulations, we note that the theoretical ICT difference aligns with the simulation results (Fig. S3, purple line). This result highlights that, as phage drive bacteria below a critical level, the immune system takes charge, effectively controlling and driving bacteria to extinction.

### F. Evaluating the impact of variations in realistic mucin levels and intermediate innate immune states on the phage therapeutic outcome

As an additional model analysis, we explore how variations in mucin and innate immune levels impact phage therapy outcomes. To model intermediate immune states, we vary the percentage of neutrophil availability in the lungs from 1% to 100%, where 100% availability corresponds to ∼3.24 × 10^6^ lung neutrophils in immunocompetent mice^32^. Then, we vary mucin levels across a physiologically relevant range for acute lung infections, ranging from 0% to 4%. Moreover, we use the phage adsorption rate, *ϕ* = 1.686 ×10^−7^ (ml/PFU)^*σ*^*h*^−1^, for our model simulations. To simulate the phage treatment of a *P. aeruginosa* infection, a host is inoculated with 10^6^ bacterial cells, and after 2 hr, the host is treated with 10^7^ phage. To assess the robustness of model predictions, we randomize the initial conditions and try 84 different ways of allocating the bacterial inoculum and the phage dose in the network. Then, we calculate the probability of clearing the infection by simulating the different initial conditions, given a specific mucin level and innate immune state.

When neutrophil availability is *>*45%, the metapopulation model predicted a ∼ 43% probability of clearing the infection regardless of mucin level (Fig. S4). The result contrasts with the prediction of the well-mixed model, where infection always clears when ≥ 45% of lung neutrophils are available (Fig. S5). Increasing neutrophil availability to 80% increases the chances of therapeutic success to 66%, especially for low mucin levels ranging from 0% to 2% (Fig. S4). The higher probability of clearing the infection in low mucin levels suggests that when metapopulation dynamics are more homogeneous, the elimination of bacteria by phage and neutrophils is facilitated. Phage and bacteria spread faster in low mucin levels, so their population dynamics homogenize among network nodes. Consequently, neutrophil resources are homogeneously distributed in the network, easing infection control. On the other hand, high mucin levels limit the diffusion of both phage and bacteria, causing them to predominantly occupy nodes proximal to their inoculation sites. This results in heterogeneous population dynamics and uneven distribution of neutrophil resources across network nodes. We hypothesize that limited neutrophil resources and high mucin levels may negatively impact the phage therapeutic outcome.

The model predicted a 100% chance of eliminating the infection when the host is fully immunocompetent (i.e., 100% neutrophil availability) and mucin levels vary between 0-3% (Fig. S4). This outcome is consistent with previous simulations, where we tested different distributions of phage dose and bacterial inoculum in a fully immunocompetent host (Fig. 4). Overall, outcomes suggest that the metapopulation spatial structure influences the dynamics of the infection and the phage therapeutic outcome.

### G. Analysis of *in vivo* imaging of *P. aeruginosa* infected mice

We use images of mice generated previously in an *in vivo* phage therapy study^13^ to extract the luminescence signal indicating the presence of a bacterial infection. In that study, authors used a *P. aeruginosa* PAKlumi strain to infect several groups of mice and the *P. aeruginosa* phage PAK P1 to treat the infected animals. They track the evolution of the infection for 72 hours, taking pictures at 2, 4, 6, 8, 24, 48, and 72 hours post-infection using the IVIS imaging system. We focus on a group of 14 mice, including immunocompetent wild-type (N=4) and lymphocyte deficient *Rag*2^−*/*−^*Il*2*rg*^−*/*−^ (N=10) mice, as they tend to survive the bacterial infection when treated with phage. Both WT and *Rag*2^−*/*−^*Il*2*rg*^−*/*−^ mice groups produce neutrophils during the bacterial infection.

The IVIS imaging system acquires a photographic image of the animal and overlays the bioluminescent infection signal on the image^55^ to create a composite picture of the infected animal. We use MATLAB R2020b and its imageprocessing capabilities to read and extract data from the time series of imaged mice. Using the images of mice, we select a region of interest (24 pixels width x 35 pixels height) that covers the animal’s lungs, throat, and nose. Then, we select a border that separates the upper and the lower compartments of the mouse respiratory system. For instance, the upper compartment includes the nose and throat, while the mouse lungs are in the lower compartment. After splitting the region of interest into two compartments, we can analyze the progression of bacterial infection in each compartment.

We pre-process the images containing bioluminescence signals and normalize the pixel intensity values of all images to a 0-1 scale. Pixel intensity 1 represents a high bacterial density, while 0 represents no signal of detected bacterial infection. We extract the infection signal from a particular compartment by adding the pixel intensity values of all the pixels that make up that compartment. We called this value the total intensity signal. We evaluate the progress of the bacterial infection in both the upper and lower compartments by examining the changes in the total intensity signal over time.

### H. Calculating infection clearance time from *in vivo* mice infection data

We are interested in calculating the time to infection resolution using the images of infected mice and comparing it with the clearance time predicted by the metapopulation model. To do so, we define an intensity threshold below which the total intensity signal is cleared (and so is the infection). By looking at the images of infected mice and calculating the total intensity of both upper and lower compartments, we set an intensity threshold of 3. When the total intensity of one compartment is above 3, we consider the infection active in that compartment.

We define some filters to consider which mice to use in the infection clearance analysis. For example, when a mouse dies before 72 hr, i.e., before the completion of the experiment, we discard that mouse for our analysis. After filtering out our dataset, we kept four immunocompetent wild-type and nine lymphocytes deficient *Rag*2^−*/*−^*Il*2*rg*^−*/*−^ mice for a total N = 13 mice.

We calculate the time to infection resolution as the time it takes for the total intensity signal to fall below the intensity threshold (conditional on the signal never exceeding the intensity threshold again after the signal falls below the threshold). When the total intensity signal of one compartment does not exceed the intensity threshold during the experiment (72 hr), we set the time to infection resolution to 0 hr for that compartment. We calculate the time to infection resolution for both upper and lower compartments. Finally, we compare the clearance time difference between the compartments.

### I. Statistical analysis

Statistical analyses were conducted in MATLAB R2020b. The one-sided Wilcoxon signed-rank test was used to compare the time to infection resolution between the upper and lower compartments of the mouse. *p <* 0.05 was considered statistically significant.

### J. Data availability

The code and data used to simulate the metapopulation model, perform the image analysis, and generate the main figures, as well as the supplementary figures, can be found in the GitHub repository at https://github.com/RogerRln/metapop_lung.

**FIG. S1:**
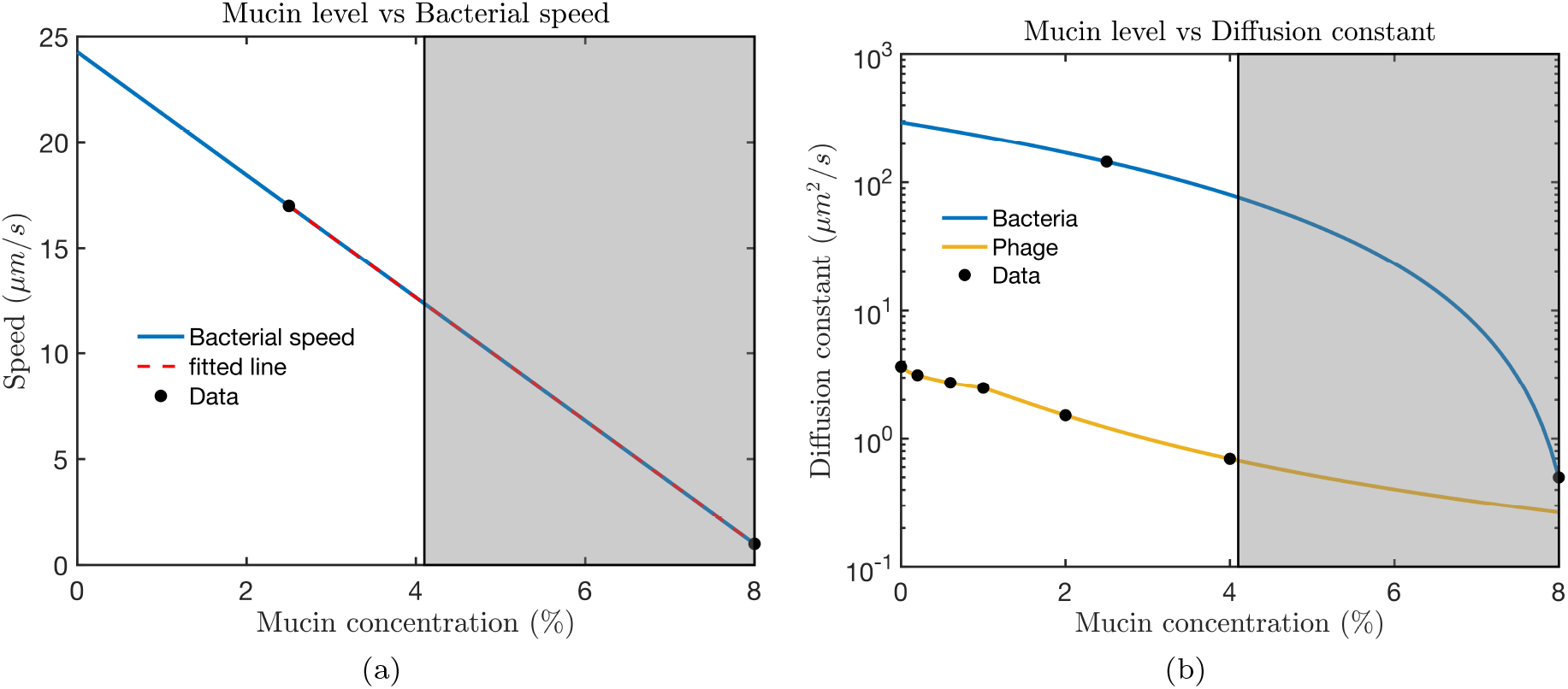
Relationship between mucin concentration and bacterial speed and phage diffusion. Bacteria speed values were collected from^29^ for two mucin levels, 2.5 and 8% (a). We fit a line (red dashed line) between the data points to find additional speed values for intermediate mucin levels (e.g., 0-4% mucin). In (b), we depict the relationship between mucin level and bacteria and phage diffusion. The gray boxes show high mucin levels not used in our model simulations.

**FIG. S2:**
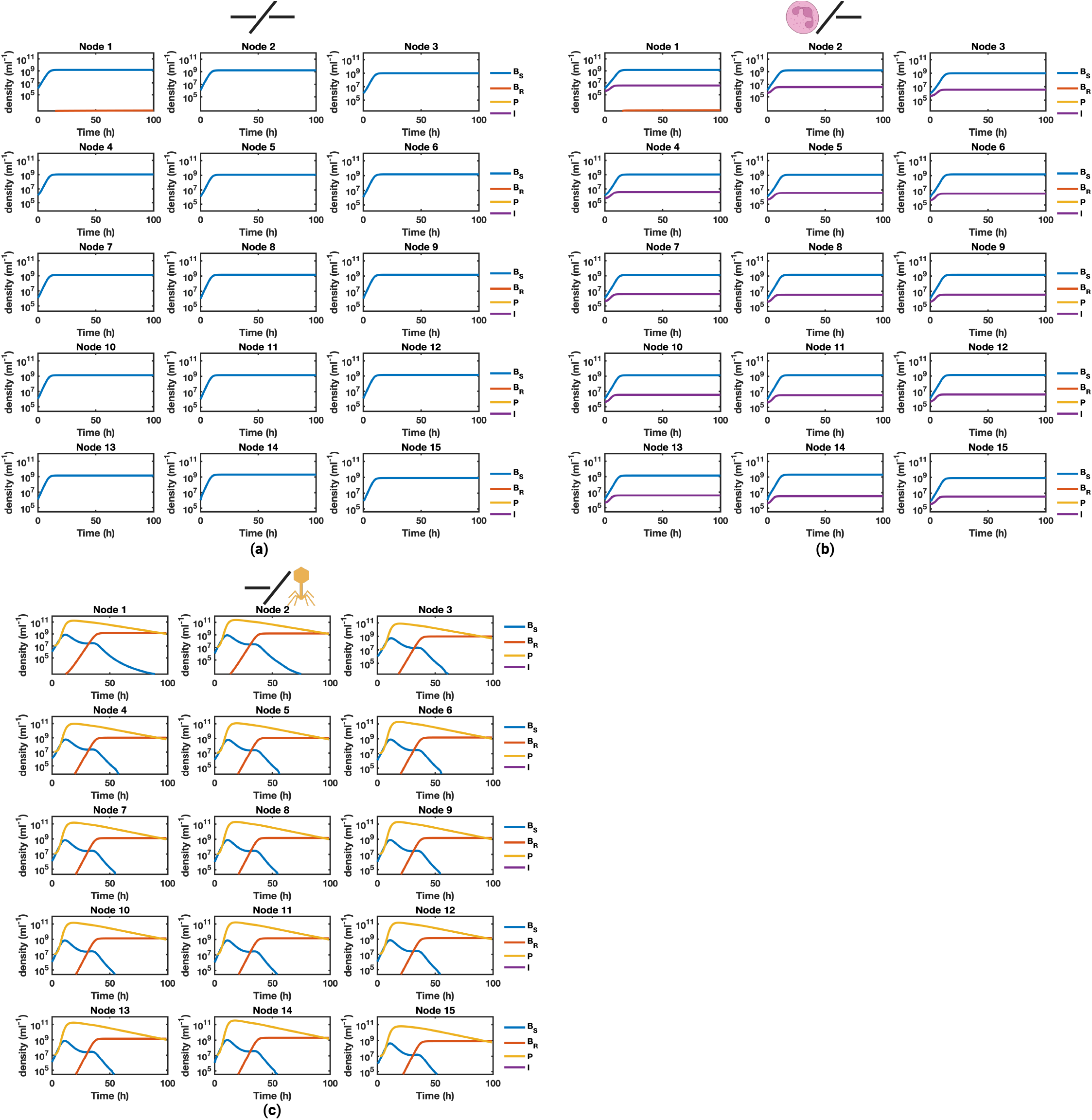
Population dynamics at the node level under different phage and immune treatments. We show the dynamics of phage (solid yellow line), phage-susceptible bacteria (solid blue line), phage-resistant bacteria (solid orange line), and the host innate immune response (purple solid line) as a result of no treatment (a), innate immune treatment (b), and phage therapy (c). We infect a host with 10^6^ bacterial cells. When used, phage (10^7^ PFU) are administered 2 hr after the bacterial infection. When the host is immunocompetent (b), we set the initial immune density to *I*_0_ = 4.05 ×10^5^ cells/ml in all the network nodes. We uniformly distribute the phage dose (c) and the bacterial inoculum (a-c) in the network such that each node had the same initial bacterial density (1.11 ×10^6^ CFU/ml) and phage density (1.11× 10^7^ PFU/ml). The simulation runs for 100 hr. Here, Node 1 = Generation 1 = trachea, and Node 15 = Generation 15 = terminal airway.

**FIG. S3:**
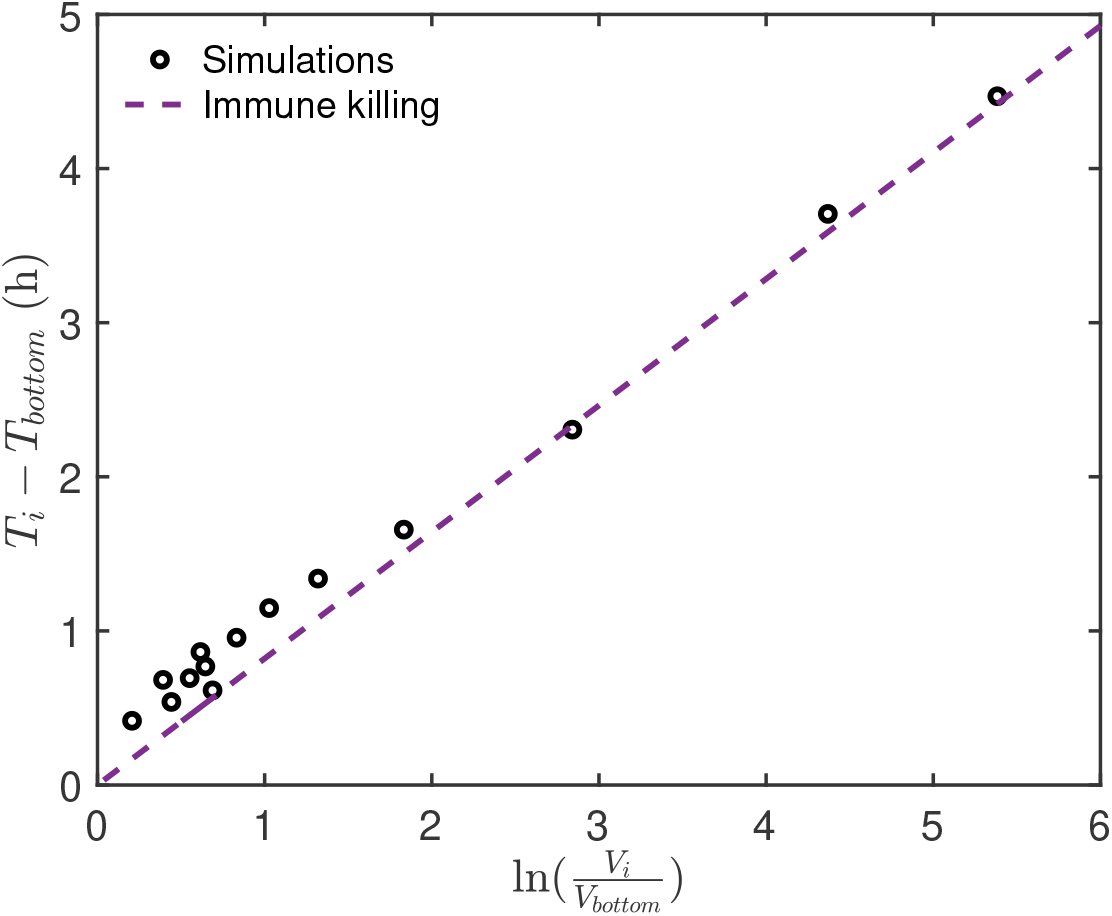
Infection clearance time difference across the network, theory vs simulations. We show the infection clearance time difference between node *i* and the last node of the network due to the immune killing rate. We compare the theoretical (dashed purple line) vs simulated (open circles) clearance time difference due to immune killing. We used the metapopulation model simulations of Fig. 2, simulating the phage therapy of a *P. aeruginosa* infection in an immunocompetent host. There, the phage dose and the bacterial inoculum were uniformly distributed such that each node had the same initial bacterial density (1.11 × 10^6^ CFU/ml) and phage density (1.11 × 10^7^ PFU/ml). The initial immune density was set to *I*_0_ = 4.05 × 10^5^ cells/ml in all the nodes.

**FIG. S4:**
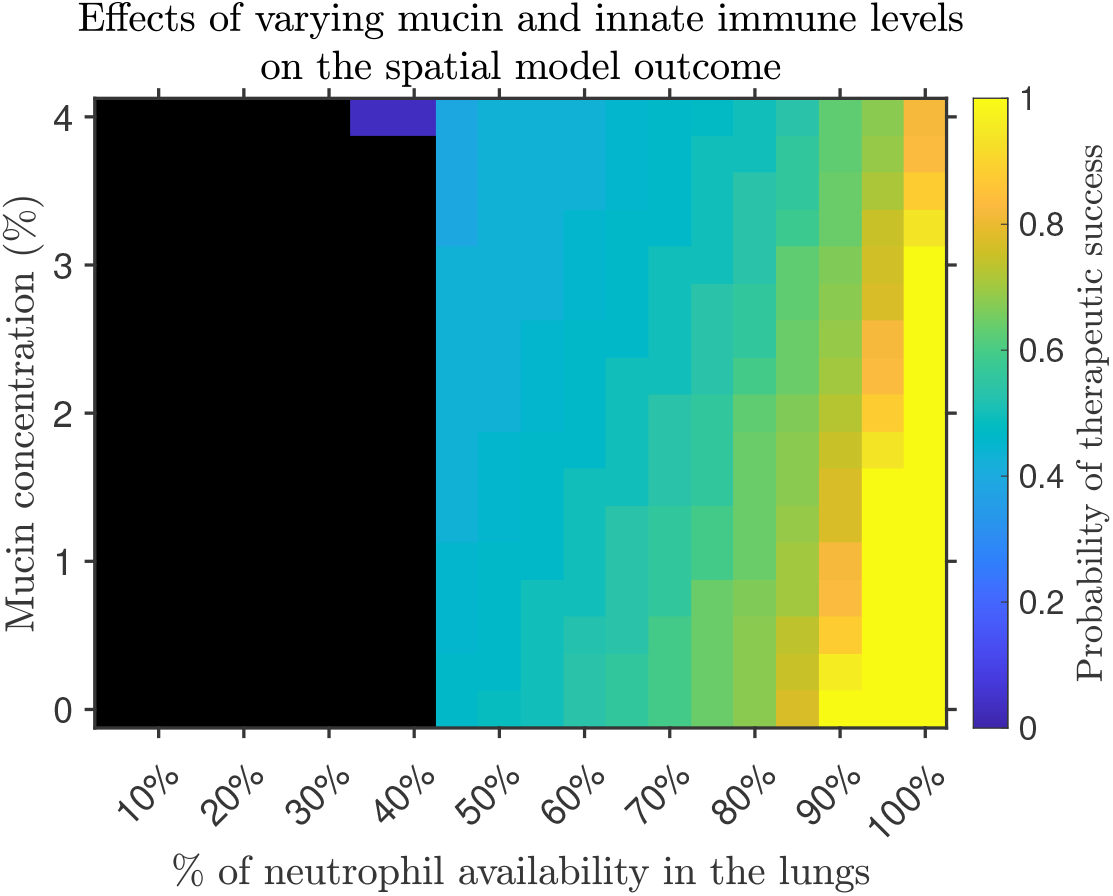
Probability of therapeutic success given intermediate mucin levels and innate immune states. To explore intermediate innate immune responses, we vary the percentage of neutrophils available in the lungs (1-100%). Further, we vary the mucin levels within a range (0-4%) that is physiologically relevant for acute lung infections. To simulate the phage treatment of a *P. aeruginosa* infection, we inoculate a host with 10^6^ bacterial cells and introduce 10^7^ phage 2 hr after the bacterial inoculation. We calculate the probability of clearing the infection by simulating 84 different initial conditions given a specific innate immune state and mucin concentration. The heatmap shows the probability of clearing the infection. The colored regions represent a *p >* 0 of clearing the infection, while black regions represent a *p* = 0 of therapeutic success. The simulation runs for 250 hr. A 100% neutrophil availability represents ∼ 3.24 × 10 lung neutrophils in an immunocompetent mouse^32^. For the simulations, we use a phage adsorption rate value of 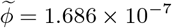 (ml/PFU)^*σ*^*h*^−1^.

**FIG. S5:**
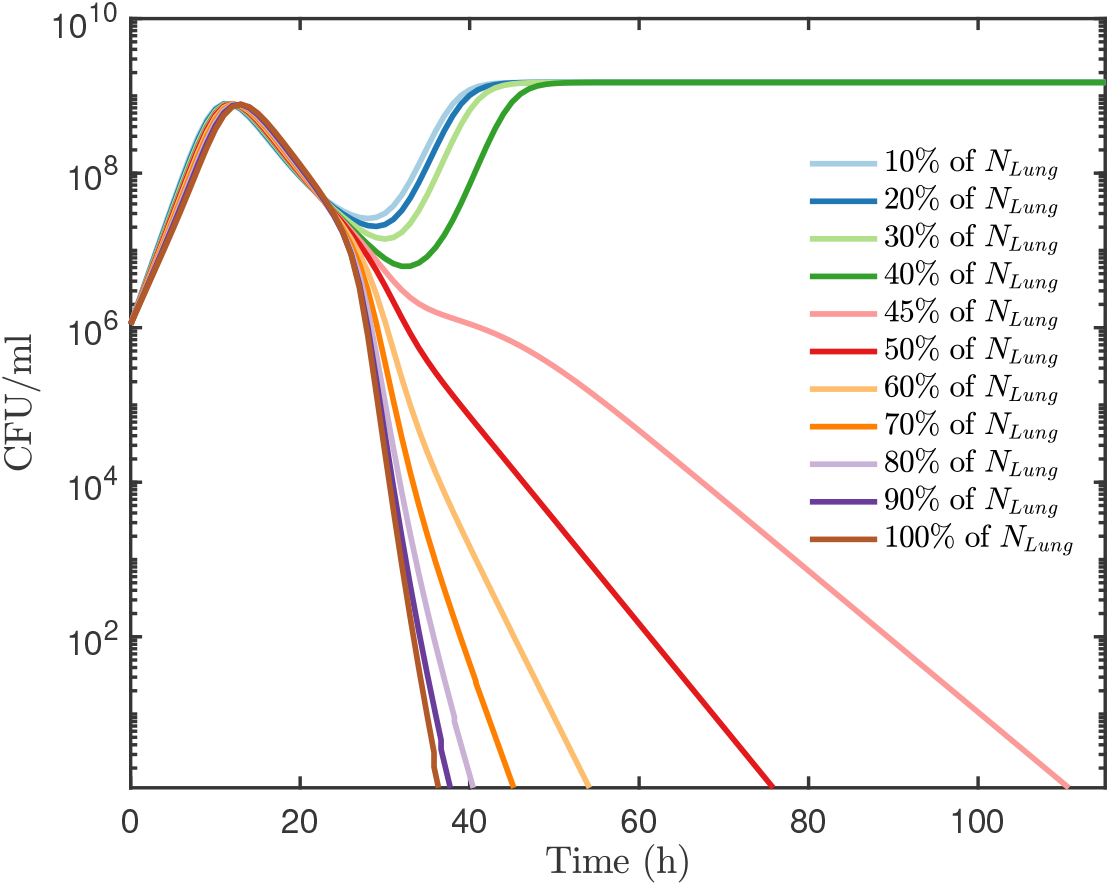
Bacterial dynamics of the well-mixed model for intermediate innate immune states. We show total bacterial dynamics (*B*_*tot*_ = *B*_*S*_ + *B*_*R*_) that result from infecting a host with 10^6^ *P. aeruginosa* cells. Phage therapy (10^7^ PFU) is administered 2 hr after the bacterial inoculation. To model intermediate immune response levels, we vary the percentage of neutrophils available in the lungs from 10 to 100%. We consider a total lung volume of 0.9 ml. The simulation runs for 115 hr. A 100% neutrophil availability represents ∼ 3.24 × 10^6^ lung neutrophils in an immunocompetent mouse^32^.

